# Feature selection followed by a novel residuals-based normalization simplifies and improves single-cell gene expression analysis

**DOI:** 10.1101/2023.03.02.530891

**Authors:** Amartya Singh, Hossein Khiabanian

**Affiliations:** Center for Systems and Computational Biology, Rutgers Cancer Institute of New Jersey, Rutgers University, New Brunswick, New Jersey; Department of Pathology and Laboratory Medicine, Rutgers Robert Wood Johnson Medical School, Rutgers University, New Brunswick, New Jersey

## Abstract

Normalization is a crucial step in the analysis of single-cell RNA-sequencing (scRNA-seq) counts data. Its principal objectives are to reduce the systematic biases primarily introduced through technical sources and to transform the data to make it more amenable for application of established statistical frameworks. In the standard workflows, normalization is followed by feature selection to identify highly variable genes (HVGs) that capture most of the biologically meaningful variation across the cells. Here, we make the case for a revised workflow by proposing a simple feature selection method and showing that we can perform feature selection before normalization by relying on observed counts. We highlight that the feature selection step can be used to not only select HVGs but to also identify stable genes. We further propose a novel variance stabilization transformation inclusive residuals-based normalization method that in fact relies on the stable genes to inform the reduction of systematic biases. We demonstrate significant improvements in downstream clustering analyses through the application of our proposed methods on biological truth-known as well as simulated counts datasets. We have implemented this novel workflow for analyzing high-throughput scRNA-seq data in an R package called Piccolo.

## Introduction

Bulk RNA sequencing (RNA-seq) studies have led to a significant improvement in our understanding of gene expression profiles associated with healthy as well as diseased states of various tissue types. However, these studies only provide an averaged view at the tissue level in which subtle but crucial distinctions of the constituent cell-types and states are obscured. Rapid advances in single-cell RNA-seq (scRNA-seq) protocols and platforms over the past decade have now facilitated investigation of transcriptional profiles at the level of individual cells, thereby enabling identification of distinct cell-types and cell states [1–5], as well as stages of development and differentiation [6, 7].

In contrast to measurements on bulk tissues, single-cell measurements have significantly greater uncertainty due to the low amounts of starting material as well as low capture efficiencies of the protocols (typically, high-throughput protocols only capture between 5% to 20% of the molecules present in each cell [8]). As a result, even deeply sequenced datasets may have up to 50% zeros [9]. The high sparsity poses a significant challenge during the computational analysis. Early attempts to build statistical models to explain the relationship between the observed counts and the true underlying gene expression levels relied on zero-inflation models to explain the excess zeros. However, data generated using newer scRNA-seq protocols that rely on unique molecular identifiers (UMIs) have been shown to be sufficiently described using simpler statistical models that do not include zero inflation [10, 11]. In this paper, any reference to scRNA-seq data will specifically pertain to UMI counts data.

A critical step in the computational analyses of both bulk RNA-seq and scRNA-seq datasets is that of normalization. The objective of normalization is to reduce the biases introduced by technical sources or even biological sources such as cell cycle state, so that we can confidently identify true biological differences [9, 12–14]. Owing to the small amount of mRNAs captured per cell, the effect of these biases is more pronounced in the case of scRNA-seq data, further underscoring the need to reduce the impact of these biases on downstream analyses. Typically, normalization is performed by re-scaling the observed counts using cell-specific size factors to reduce the differences in sampling depths (total counts) between the cells. The scaled counts are then transformed with the help of a monotonic non-linear function (usually the logarithm function) to stabilize the variances of genes across different mean expression levels (such a transformation is popularly referred to as a variance stabilization transformation).

In the standard scRNA-seq workflow (implemented for instance in Seurat [15–18] and Scanpy [19]) normalization is followed by a feature selection step that focuses on identifying genes that capture most of the biological variation across the cells while eliminating genes that do not exhibit meaningful biological variation. This sequence of steps - normalization followed by feature selection - in the standard workflow appears quite reasonable, especially given the fact that differentially expressed genes can be identified reliably only after reducing the sampling depth differences between the cells. However, objective (i). identification of genes that are differentially expressed between groups of cells, is not the same as objective (ii). identification of highly variable genes (HVGs). While the identification of differentially expressed genes between distinct groups of cells requires that the sampling depth differences be reduced through normalization, we show that it isn’t necessary to perform normalization in order to identify HVGs.

In this article, we first revisit and re-examine the fundamental nature of the scRNA-seq counts data obtained from high-throughput technologies. Aided by a clearer understanding of the nature of the data, we propose a simple feature selection method that relies on a regression-based approach to estimate dispersion coefficients for the genes based on their observed counts. Using this method, we show that feature selection can be reliably performed before normalization. Importantly, during the feature selection step we not only identify variable genes, but also shortlist stable genes. The variation in the counts of these stable genes is expected to primarily reflect the biases introduced by the technical sources, and can therefore be used to estimate cell-specific size factors in order to perform normalization. During normalization we also need to ensure variance stabilization, especially when relying on principal components analysis (PCA) for dimensionality reduction. Keeping this in mind, we propose a residuals-based normalization method that not only reduces the impact of sampling depth differences between the cells but simultaneously ensures variance stabilization by explicitly relying on a monotonic non-linear transformation (default choice is the *log* transformation). We demonstrate significant improvements in downstream clustering analyses enabled by the application of our feature selection and normalization methods on biological truth-known as well as simulated counts datasets. Based on these results, we make the case for a revised scRNA-seq analysis workflow in which we first perform feature selection and subsequently perform normalization using our residuals-based approach. We have implemented this novel scRNA-seq workflow in an R package called `Piccolo`.

## Results

### Genes with small counts exhibit quasi-Poisson variance

#### Genes with low mean expression levels show Poisson-like variance of their counts

As pointed out by Sarkar and Stephens [11], a good starting point for understanding the nature of the scRNA-seq counts data is to recall that the observed counts reflect contributions from both the underlying expression levels of the genes as well as the measurement errors. This necessitates that the contributions from the two be carefully distinguished in order to better understand and explain the true biological variation. Based on their analysis, they found that a simple Poisson distribution sufficed to explain measurement error, while a simple Gamma distribution often sufficed to explain the variation in expression levels across cells. The observation model built using these two distributions yields the Gamma-Poisson (or negative binomial (NB)) distribution which is well-known as a plausible model to explain over-dispersed counts. Under this model, the mean-variance relationship is given by,

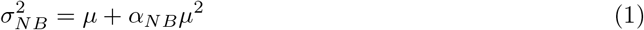

where *α*_*NB*_ is the NB over-dispersion coefficient (*α*_*NB*_ = 0 yields the familiar Poisson mean-variance relationship: *σ*^2^ = *μ*).

A more familiar expression for the NB mean-variance relationship is,

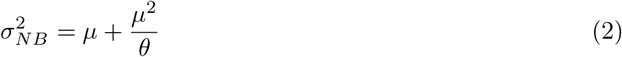

where *θ* is referred to as the inverse over-dispersion coefficient.

Another mean-variance relationship closely related to the Poisson and the NB is the quasi-Poisson (QP) wherein,

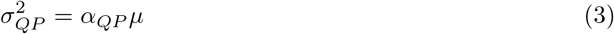

where *α*_*QP*_ is the QP dispersion coefficient (*α*_*QP*_ = 1 yields the familiar Poisson mean-variance relationship; *α*_*QP*_ *>* 1 would be associated with counts over-dispersed with respect to the Poisson, while *α*_*QP*_ *<* 1 would indicate counts under-dispersed with respect to the Poisson).

We began our investigation into the nature of UMI counts by considering the mean-variance relationship of counts for genes in a technical negative control data set [10](hereafter referred to as `Svensson 1`). This data set consisted of droplets containing homogenous solutions of endogenous RNA as well as spike-in transcripts. The variability of counts in this data arises solely due to technical sources and is not attributable to any underlying biological source. The left panel in Fig. 1A shows the variance (*σ*^2^) vs mean (*μ*) log-log plot for `Svensson 1`. Each dot in the plot corresponds to a gene. The colors of the dots reflect the local point density, with darker shades (deep blues) indicating low density and brighter shades (bright yellow) indicating high density. The black line depicts the variance expected under the Poisson model (*σ*^2^ = *μ*).

**Figure 1.**
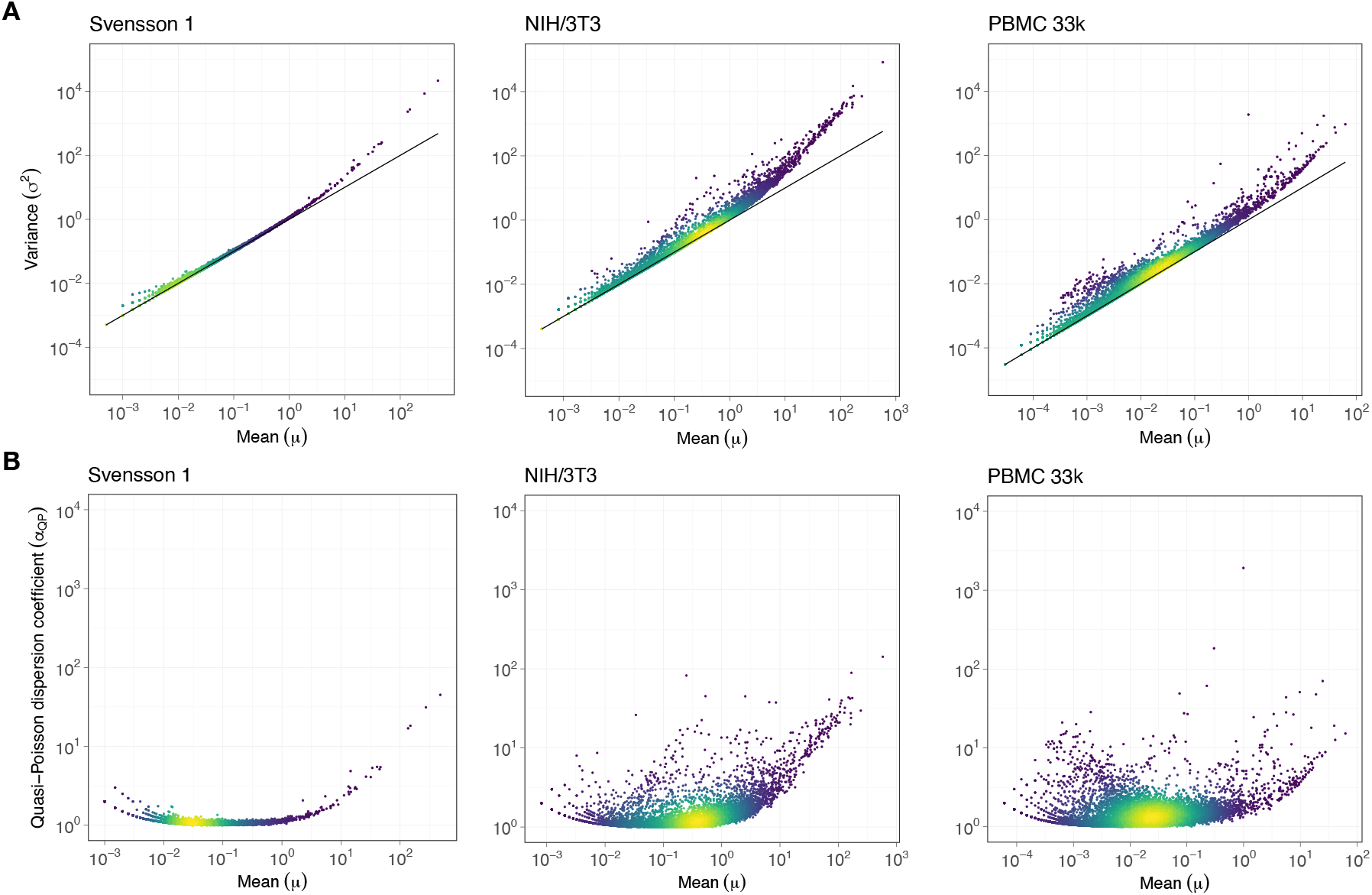
Genes with low mean expression exhibit quasi-Poisson variance. In all the plots, each dot represents a gene and the color of the dots reflect the local point density, with brighter shades (yellow) indicating high density and darker shades (deep blue) indicating low density. **A**. Variance (*σ*^2^) vs mean (*μ*) log-log scatter plots for the Svensson 1 technical control (left panel), NIH/3T3 fibroblast cell line (center panel), and PBMC 33k (right panel) datasets. The solid black line corresponds to the Poisson model (*σ*^2^ = *μ*). For genes with low mean expression levels, the variance can be adequately described by the Poisson model. **B**. Quasi-Poisson dispersion coefficients (*α*_*QP*_) vs mean (*μ*) log-log scatter plots for the Svensson 1 (left panel), NIH/3T3 (center panel), and PBMC 33k (right panel) datasets. *α*_*QP*_ for each gene were estimated from the observed counts using a regression-based approach.

We begin by noting that the UMI counts are heteroskedastic in nature since the variances of the genes depend on the mean (larger variances corresponding to larger means). If we then move on to focus on genes with low mean expression levels (especially *μ <* 0.1), we can see that the mean-variance relationship appears to be well-approximated by the Poisson model since their observed variances lie close to the black line. It’s only for genes with higher mean expression levels (especially *μ >* 1) that the variance of the observed counts exceed the variance expected under the Poisson model. For comparison with counts data with inherent biological variation, we looked at the NIH/3T3 fibroblast cell line data set [10] (hereafter referred to as `NIH/3T3`) and the 10X Genomics Peripheral Blood Mononuclear Cells (PBMC) 33k data set [20] (hereafter referred to as `PBMC 33k`). Even for these two datasets, most of the genes with low mean expression levels exhibit Poisson-like mean-variance relationship (middle and right panels in Fig. 1A).

The Poisson-like nature of the mean-variance relationship at low expression levels can be made clearer with the help of a simple example. Consider a gene with *μ* = 0.01. Let us assume that the counts of this gene are actually NB distributed and that 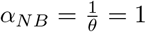 is the true estimate of the over-dispersion coefficient. The variance of the counts will then be 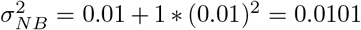. In comparison, the expected variance under the Poisson model will be 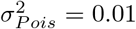. The percent difference between the two variances is a mere 1%. Thus, even if we approximated the variance for the counts of this gene with that predicted by the simple Poisson model, the estimated variance would differ by only 1% from the true NB variance. The percentage difference between the Poisson and the NB variances would of course increase for larger values of *α*_*NB*_, however, estimates of *α*_*NB*_ for real biological datasets typically range between 0.01 and 1 [21, 22].

We can formalize the discussion above with the help of equations. 1 and 3. We note that when *α*_*NB*_*μ <<* 1,

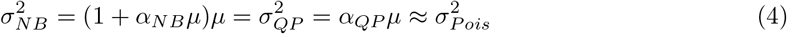

where we defined *α*_*QP*_ in terms of *α*_*NB*_ and *μ* as *α*_*QP*_ = 1 + *α*_*NB*_*μ* (see Appendix for a discussion on viewing QP variance as a special case of NB variance).

Thus, we conclude that counts of genes with low mean expression levels and moderate over-dispersion (such that *α*_*NB*_*μ <<* 1) exhibit variances that do not deviate significantly from the variances predicted under the Poisson model.

#### QP dispersion coefficients can be obtained using a regression-based approach

For the NB distribution, the usual approach is to use maximum-likelihood estimation (MLE) to obtain estimates for the over-dispersion parameter (*α*_*NB*_). This approach apart from being computationally intensive has the weakness that if in fact the distribution is not NB, the maximum-likelihood estimator is inconsistent. Cameron and Trivedi proposed a regression-based test that offers a more robust alternative by requiring only the estimates of the mean and variance for each gene [23, 24]. The test is set up to estimate over-dispersion beyond the null model (Poisson distribution) by specifying the alternate model in the form of a scalar multiple of a function of the mean,

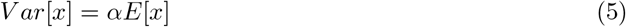

where the scalar multiple (*α*) is estimated using least squares regression (see Appendix).

The QP mean-variance relation (see equation. 3) corresponds precisely to such an alternative hypothesis in which the variance is simply a scalar multiple (*α*_*QP*_) of the mean. This enables us to use this simple yet robust regression-based approach to estimate the QP dispersion coefficients (*α*_*QP*_) for each gene by simply relying on the estimates of their mean and variance.

#### QP variance for counts of genes with low mean expression is simply due to the observed counts being small and is not biological in origin

We obtained estimates for *α*_*QP*_ for all genes using the regression-based approach. Based on these estimates, we filtered out genes that were under-dispersed compared to the Poisson model (*α*_*QP*_ *<* 1). For the remaining genes, we plotted *α*_*QP*_ vs *μ* log-log plots (Fig. 1B). For `Svensson 1` (left panel in Fig. 1B), it is evident that for genes with low mean expression levels their *α*_*QP*_ values lie close to 1 (10^0^). In particular, we note that the mean of the *α*_*QP*_ for genes with *μ <* 0.1 is 1.0474. This shows that for these genes with low mean expression their *α*_*QP*_ values are close to 1 which is consistent with the observation that the counts of these genes exhibit Poisson-like variance. Since `Svensson 1` is a technical control data set, this observation further supports the simple Poisson distribution as an appropriate model for explaining measurement error, particularly for genes with low mean expression levels. Furthermore, genes with *μ <* 0.1 do not appear to exhibit any dependence on *μ*. We evaluate this quantitatively by using the non-parametric Kendall’s rank and Spearman’s rank correlation tests to determine whether there is a statistical dependence between *α*_*QP*_ and *μ* values for genes with *μ <* 0.1 (see Methods). Both tests evaluate how well the relationship between two variables can be described using a monotonic function. For both tests, the correlation coefficients - *τ* and *ρ* respectively - indicate a statistical dependence if the values are close to +1 or − 1, while values of *τ* or *ρ* closer to 0 indicate the absence of such a statistical dependence. The resultant correlation coefficient values of *τ* = 0.04611706 (*p* = 0.001244) and *ρ* = 0.0880 (*p* = 3.475*E* − 05), support the assertion that there is no statistical dependence between *α*_*QP*_ and *μ* values for genes with low mean expression levels.

For `NIH/3T3` (middle panel in Fig. 1B) and `PBMC 33k` (right panel in Fig. 1B), despite greater variability in *α*_*QP*_ due to the inherent biological variability in the data, similar observations were made - namely that the *α*_*QP*_ values are close to 1 for genes with *μ <* 0.1 and that there was a lack of dependence between *α*_*QP*_ and *μ* for those genes. For `NIH/3T3`, the mean of the *α*_*QP*_ for genes with *μ <* 0.1 is 1.1388, and both Kendall’s correlation coefficient *τ* = 0.0644 (*p* = 1.28*E* − 05) and Spearman’s correlation coefficient *ρ* = 0.1014 (*p* = 4.084*e* − 06) indicate that there is no statistical dependence between *α*_*QP*_ and *μ* for genes with *μ <* 0.1. For `PBMC 33k`, the mean of the *α*_*QP*_ for genes with *μ <* 0.1 is 1.3710, and the Kendall’s correlation coefficient *τ* = 0.1349 (*p <* 2.2*E* − 16) and Spearman’s correlation coefficient *ρ* = 0.2035 (*p <* 2.2*E* − 16) once again suggesting that there is no significant statistical dependence between the *α*_*QP*_ values and *μ* for genes with low mean expression levels.

To summarize, we observed that the counts for genes with low mean expression levels exhibit QP variance (see Appendix for a discussion on how the lack of dependence between *α*_*QP*_ and *μ* for genes with low mean expression levels manifests as a non-decreasing relationship between their *θ* and *μ*). It is important to highlight here that we observed this relationship not just in the biological datasets (`NIH/3T3` and `PBMC 33k`) but also in the technical control data set (`Svensson 1`), suggesting that this relationship is not biological in origin and can be understood more simply in terms of the fact that for genes with small counts the variance of those counts can barely exceed the variance expected under the Poisson distribution.

### Feature selection can be performed before normalization

#### Standard scRNA-seq workflow is based on an assumption that reflects a confusion between the distinct objectives of identification of variable genes and differentially expressed genes

In the standard scRNA-seq workflow, identification of HVGs - genes that exhibit greater variability in counts compared to other features in the data set - is performed only after normalization. This particular sequence in the workflow is based on the assumption that unless the systematic biases are reduced or eliminated, we cannot reliably identify features that best capture the biological variability inherent in the data. A more careful examination reveals that this assumption reflects a confusion between two fundamentally distinct objectives. The first objective - identification of HVGs - as mentioned above, is about identifying genes that exhibit higher variability of counts compared to other features in the data set; we expect that these genes capture most of the biological variability in the data set. The second objective - identification of differentially expressed (DE) genes - is about identifying genes that exhibit differences in their expression levels between distinct sets of cells. For identifying DE genes, it is indeed imperative that the systematic biases are reduced or eliminated in order to identify genes that truly reflect actual biological differences between the distinct groups of cells being compared. However, it is possible to identify HVGs without first accounting for the systematic biases through normalization since these biases owing to their systematic nature are expected to manifest as additional but *consistent* sources of variation for the counts of genes across cells. This is in fact the assumption underlying the normalization approaches that rely on estimation of cell-specific size factors to re-scale and adjust the observed counts.

We can state the expectation discussed above in terms of the QP dispersion coefficients. Suppose we have gene *A* and gene *B* with comparable mean expression levels (*μ*_*A*_ ≈ *μ*_*B*_) such that,

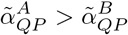

where 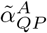 and 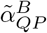 are the QP dispersion coefficients of gene A and gene B in the hypothetical case where there is no systematic bias. Our expectation is that even in the presence of systematic biases, the relative magnitudes of the dispersion coefficients for the bias-affected counts will be such that,

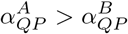

where 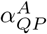 and 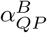 are the respective QP dispersion coefficients obtained from the observed counts of genes *A* and *B*. Thus, the expectation is that genes that exhibit high variability in their counts due to underlying biological differences will exhibit high variability even in the presence of systematic biases. We can test whether this is a reasonable expectation by introducing systematic biases into a given data set and then verifying whether the genes we identify as variable for the modified data set are consistent with the ones identified for the original data set. Before proceeding to perform such a test, however, we first need to lay down a method to identify HVGs.

#### Feature selection method based on QP dispersion coefficients identifies HVGs and stable genes

As discussed earlier, distinguishing between measurement and biological processes by separately modeling their contributions to observed counts enables a better understanding and interpretation of the observed counts [11]. It also provides a simple and straightforward basis for identifying and filtering out features that are relatively uninformative prior to any downstream analysis: if the measurement model sufficiently describes the observed counts for a given gene, then no meaningful biological inference can be drawn based on the counts of that gene. This is so since all the variation in the counts for that gene can be attributed to the measurement process itself, with negligible or no contribution from any biological process. We call such genes whose counts are adequately described by the measurement model as *biologically uninformative*. Since a simple Poisson distribution suffices to explain measurement error, using the estimates for *α*_*QP*_ we can easily identify genes that are likely to be biologically uninformative - genes with *α*_*QP*_ ≤ 1 do not exhibit over-dispersion with respect to the Poisson measurement model and can be filtered out.

While we can simply rely on the *α*_*QP*_ to identify biologically uninformative genes, identification of HVGs based on just the magnitudes of *α*_*QP*_ would result in a bias towards genes with higher mean expression levels (see Fig. 1B). In order to address this and ensure that there is no preferential selection of genes with higher mean expression levels, we propose the following approach to shortlist HVGs:

- Group the genes into bins (default choice is 1000 bins) based on their mean expression levels; each bin contains approximately the same number of genes with comparable mean expression levels
- Sort the genes within each bin into quantiles based on their *α*_*QP*_
- Obtain the *α*_*QP*_ corresponding to the reference quantile within each bin (default reference quantile is the 10th quantile) - we refer to this as *α*_*QP* (*Reference*|*Bin*)_
- Calculate *α*_*QP*_ − *α*_*QP* (*Reference*|*Bin*)_ for each gene - larger values indicate greater over-dispersion

We illustrate the binning process involved in our feature selection method for `Svensson 1` (Fig. 2A). For ease of illustration we show 10 bins (see Appendix and Fig. S6 for a discussion on how the number of bins impact the identification of HVGs), where each bin corresponds to a segment or region between the vertical dashed lines. Since `Svensson 1` is a technical control data set, we don’t expect to see much variability in the expression levels of the genes, and indeed in Fig. 2C we can see that there are just two genes (spike-in transcripts ERCC-00074 and ERCC-00130) that exceed the threshold of 20 for *α*_*QP*_ − *α*_*QP* (*Reference*|*Bin*)_ (red horizontal dashed line). For `PBMC 33k`, in contrast, we observe 15 genes that exceed the threshold of 20 for *α*_*QP*_− *α*_*QP* (*Reference*|*Bin*)_ (Fig. 2D). The threshold of 20 was picked simply to illustrate how the HVGs may be shortlisted. In practice, we can rank the genes based on the magnitudes of *α*_*QP*_− *α*_*QP* (*Reference*|*Bin*)_ and shortlist the top 3000 genes. An important point to highlight here is that the shortlisted HVGs do not exclude genes with low mean expression - note the inset panel in the top left corner of Fig. 2D, where we show the histogram based on the mean expression levels for the HVGs (darker shade) together with the histogram for all genes with *α*_*QP*_ *>* 1 (lighter shade).

**Figure 2.**
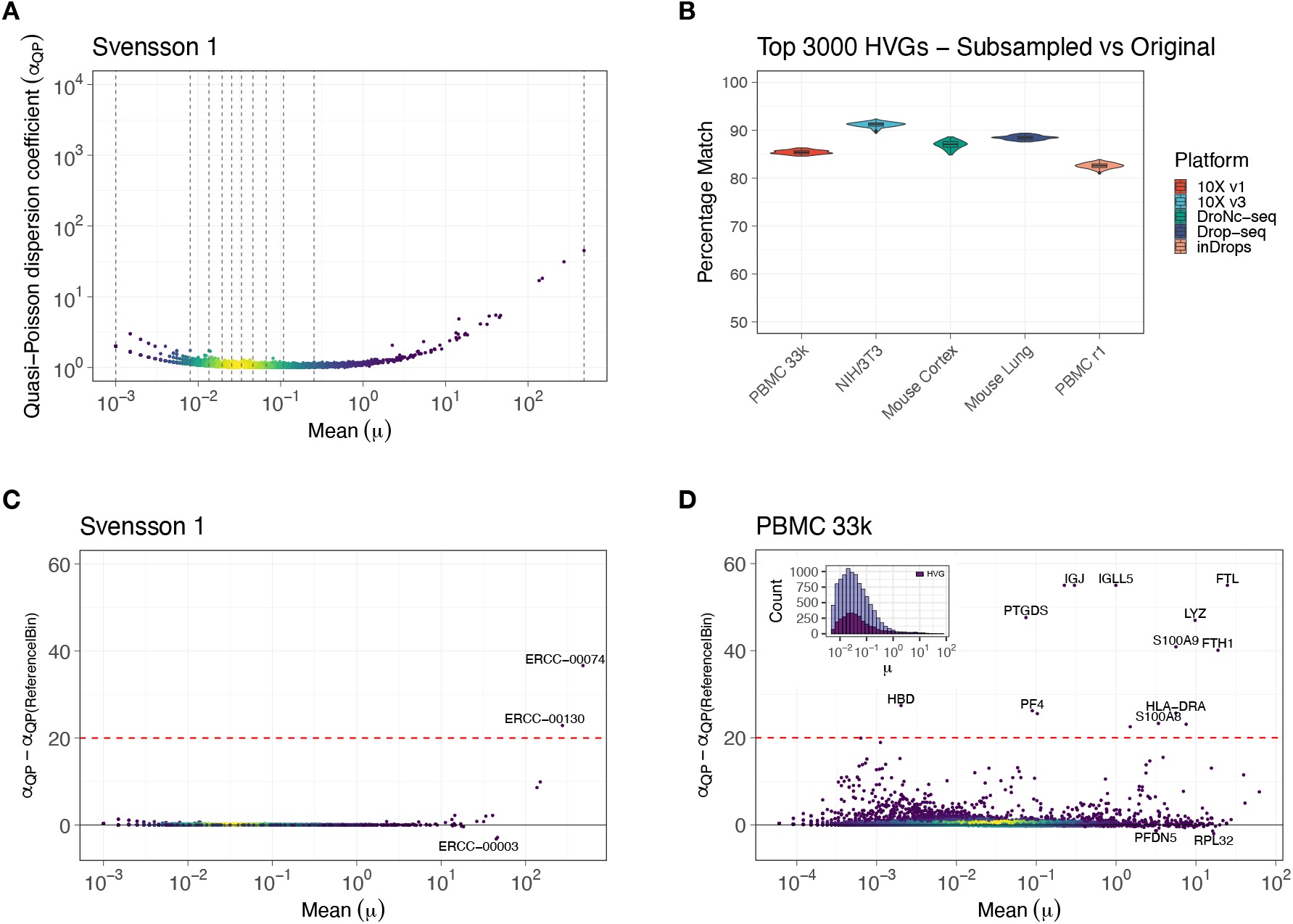
Feature selection can be performed before normalization based on dispersion coefficients estimated using the observed counts. **A**. Quasi-Poisson dispersion coefficient (*α*_*QP*_) vs mean (*μ*) log-log scatter plots for genes that exhibited over-dispersion with respect to the Poisson (*σ*^2^ *> μ*) for the Svensson 1 technical control data set. Each dot represents a gene. The dashed gray vertical lines illustrate how the genes are binned based on their mean expression levels (in this figure, there are 10 bins). Genes within adjacent pairs of dashed vertical lines belong to the same bin. For each bin, the default choice of *α*_*QP* (*Reference*|*Bin*)_ is the *α*_*QP*_ corresponding to 10th quantile within the bin. **B**. Violin-box plots showing the percentage match between the top 3000 highly variable genes (HVGs) identified by our feature selection method for the original (unsubsampled) and corresponding 100 subsampled datasets. The original 5 datasets (PBMC 33k, NIH/3T3 cell line, Mouse cortex, Mouse lung, and PBMC r1) were obtained using different single-cell platforms - 10X Chromium v1, 10X Chromium v3, DroNC-seq, Drop-seq, and inDrops respectively. The box/violin plots are colored according to the platforms. For all 5 datasets, more than 80% of the top 3000 HVGs shortlisted for the subsampled datasets matched the top 3000 HVGs of the original datasets. **C**. *α*_*QP*_ − *α*_*QP* (*Reference*|*Bin*)_ vs mean (*μ*) linear-log scatter plot for genes that exhibited over-dispersion with respect to the Poisson (*σ*^2^ *> μ*) for the Svensson 1 data set. The dashed red horizontal line illustrates the threshold to shortlist HVGs. Genes with *α*_*QP*_ − *α*_*QP* (*Reference*|*Bin*)_ *>* 20 in this case are shortlisted as HVGs. On the other hand, genes with *α*_*QP*_ − *α*_*QP* (*Reference*|*Bin*)_ *<* 0 are shortlisted as stable genes. **D**. *α*_*QP*_ − *α*_*QP* (*Reference*|*Bin*)_ vs mean (*μ*) linear-log scatter plot for genes that exhibited over-dispersion with respect to the Poisson (*σ*^2^ *> μ*) for the PBMC 33k data set. Labeled genes with *α*_*QP*_ − *α*_*QP* (*Reference*|*Bin*)_ *>* 20 are shortlisted as HVGs. On the other hand, genes with *α*_*QP*_ − *α*_*QP* (*Reference*|*Bin*)_ *<* 0 are shortlisted as stable genes (some labeled in figure). Inset in top left corner - light colored histogram corresponding to all genes with *α*_*QP*_ *>* 1, and dark colored histogram corresponding to the top 3000 HVGs. Note that the HVGs are shortlisted across the different expression levels with no preferential selection based on the mean expression level of the genes.

Aside from identifying variable genes, we can also shortlist genes that do not exhibit much variability in their counts. We refer to these genes as *stable genes*. Identification of such genes is very useful since we expect that the variability in the counts of these genes is primarily attributable to the measurement process and is not confounded by biological differences between the cells. Thus, we can rely on these genes to obtain more reliable estimates for the cell-specific size factors in order to reduce the impact of the differences in sampling depths. Based on our feature selection method, genes with *α*_*QP*_ − *α*_*QP* (*Reference*|*Bin*)_ *<* 0 are shortlisted as stable genes. For `PBMC 33k`, two of these genes (*PFDN2* and *RPL32*) are labeled in Fig. 2D below the horizontal black line at *α*_*QP*_ − *α*_*QP* (*Reference*|*Bin*)_ = 0.

### HVGs can be consistently identified despite the introduction of systematic biases

Having introduced the feature selection method, we can now test the expectation that the HVGs can be identified despite the presence of systematic biases. To perform the test, we picked 5 UMI counts datasets obtained from different platforms: `PBMC 33k` (10X Genomics Chromium v1), `NIH/3T3` (10X Genomics Chromium v3), `Mouse Cortex` r1 (DroNC-seq) [8], `Mouse Lung` (Drop-seq) [25], and `PBMC r1` (inDrops) [8]; r1 denotes replicate 1. For each of these datasets, we randomly picked 30% of the cells and subsampled the counts in those cells to a fraction of the original total count. The fractions were allowed to take one of the following values - 0.3, 0.4, 0.5, 0.6, 0.7 - and were picked randomly for each cell. We did this 100 times for each of the 5 datasets. Using our feature selection method, we shortlisted the top 3000 HVGs in each of the subsampled datasets and compared with the top 3000 HVGs in the respective original (unsubsampled) datasets to see how many of the top 3000 HVGs matched between the two. We observed that on average more than 80% of the top 3000 HVGs shortlisted for the subsampled datasets matched the top 3000 HVGs obtained from the original datasets across the different platforms and tissue types (Fig. 2B). Thus, despite the introduction of random systematic biases there is good agreement between the HVGs obtained for the original datasets and the HVGs obtained for corresponding datasets with the introduced biases. This supports our assertion that feature selection can be performed prior to normalization.

### Normalization can be performed using a residuals-based approach that includes variance stabilization

#### Cell-specific size factors should be estimated using stable genes

The basic objective of normalization is to reduce systematic biases introduced due to technical or potentially uninteresting biological sources (such as cell size, cell cycle state) before further downstream analyses such as clustering and differential expression. The most common approach to reduce such systematic biases is to scale the counts within each cell by cell-specific *size factors* [13, 26]. *The simplest estimates are given by*,

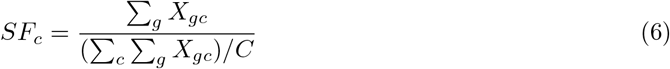

where *X*_*gc*_ is the observed count of gene *g* in cell *c* (*N*_*c*_), and *C* is the total number of cells. The numerator in equation. (6) is the total UMI count in cell *c*, while the denominator is the mean of the total counts of the cells.

There is an intimate link between the estimates of size factors given by equation. (6) and the estimates for expected means 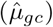 under the assumption that the counts are Poisson distributed should be viewed as estimates under the approximation that the counts are Poisson distributed (see Appendix), which suggests that the size factor estimates obtained from equation. (6). Keeping this in mind, we argue that it is most appropriate to calculate the size factors by relying on the stable genes identified using our feature selection method. The counts of these genes do not exhibit significant over-dispersion compared to the Poisson, and as discussed earlier, we expect that the primary source of the variability of their counts is the measurement process which is what we are trying to account for with the help of the size factors. Therefore, we propose the following refinement to the estimation of the size factors,

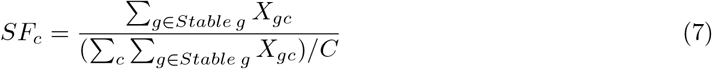

where *Stable g* refers to the set of stable genes.

#### Limitations of widely used normalization methods

In the standard scRNA-seq workflow, normalization and feature selection are typically followed by dimensionality reduction using principal components analysis (PCA). An important step prior to PCA is to ensure variance stabilization by transforming the size factor adjusted counts using the *asinh, log, sqrt* functions etc. This ensures that genes with higher expression levels (and as a consequence larger variance) do not contribute disproportionately to the overall variance as evaluated through PCA. The *log*-based variance stabilization transformation (hereafter referred to as `logSF`) is the most popular method according to which the transformed counts are given by,

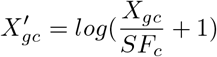

here the pseudocount of 1 ensures that the *log* transformation works with zero counts, and in fact returns zeros for these counts even after transformation. However, the `logSF` normalization is not very effective since the total counts of the cells (*N*_*c*_) shows up as a primary source of variation in PCA even after normalization (see Fig. S8). This can be traced to the fact that under this transformation, the zeros remain zeros while only the non-zero counts are scaled according to the size factors. Systematic differences in the number of zero counts between the cells can therefore be identified as a major source of variation even after transformation [27, 28].

Residuals-based approaches proposed by Hafemeister et al. [20], Townes et al. [27], and more recently by Lause et al. [21] provide alternatives that lead to much more effective normalization (see Fig. S9). The Pearson residuals under the NB model are approximately given by,

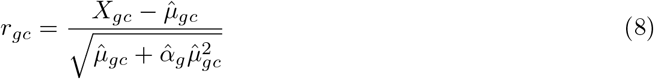

where *X*_*gc*_ is the observed count of gene *g* in cell *c*, 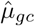 is the estimated mean of gene *g* in cell *c*, and 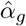 is the estimated over-dispersion parameter for gene *g*.

The increased effectiveness of normalization with the Pearson residuals can be attributed to 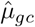 taking into account the systematic differences in the total counts between the cells. Unlike the case with `logSF` normalization where zero counts are transformed back to zeros, the zero counts are instead transformed to negative residual values whose magnitudes vary depending on the total counts of the respective cells.

The rationale underlying the residuals-based approach is that the null model should correspond to the measurement process so that the residuals provide estimates for deviations away from the expectations under the measurement model. However, the sparsity and the skewed nature of the counts distributions pose significant challenges to achieving effective variance stabilization with the help of residuals-based methods. The lack of variance stabilization becomes especially noticeable for genes that are robustly expressed in only a subset of cells while showing negligible expression in the rest of the cells (such genes would be considered as *markers* of the specific cell sub-populations in which they are expressed) [22] (see Appendix for more discussion on this). Furthermore, for genes with very low mean expression levels (and as a consequence extremely small estimated standard deviations) even cells with just 1 or 2 UMI counts sometimes end up with unusually large residual values that are then addressed through heuristic approaches [20, 21, 29].

#### Residuals-based normalization that includes variance stabilization

Keeping in mind the limitations of the widely used normalization methods discussed above, we propose a conceptually simple residuals-based normalization method that reduces the influence of systematic biases by relying on size factors estimated using stable genes while simultaneously ensuring variance stabilization by explicitly relying on a variance stabilization transformation.

In order to motivate our approach, we begin by pointing out that *z*-scores are simply Pearson residuals corresponding to the normal distribution. Assuming counts, *Y*_*gc*_, that are normally distributed,

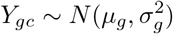

the MLEs for *μ*_*g*_ and 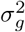 are given by, 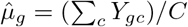 and 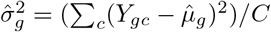, respectively. The corresponding Pearson residuals-based on these estimates are given by,

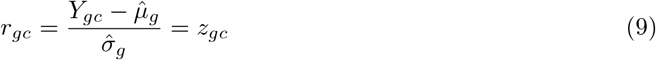

which as already noted above correspond to *z*-scores (for simplicity, we assumed that there are no differences in total counts between the cells).

In order to compute *z*-scores for our data, we first need to apply a variance stabilization transformation to the raw counts (*X*_*gc*_) to bring their distribution closer to the normal distribution. The variance stabilization transformation can be performed using monotonic non-linear functions, *g*(*X*), such that the transformed counts are given by,

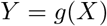

To compute the residuals, we need estimates for the means and variances of *Y* based on estimates for means and variances of *X*. We can arrive at approximations for both using a Taylor series expansion around *X* = *μ* (see Appendix). In particular for *g*(*X*) = *log*(*X* + 1), the first order approximations of the mean and variance are given by (the first-order approximation to the variance of a transformed random variable is also known as the Delta method attributed to R. A. Dorfman [30]),

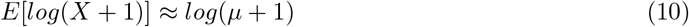

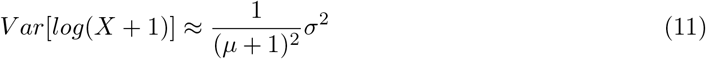

At this point we need estimates for the means 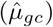 and the variances 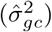 that account for the systematic biases. Note, that the usual estimate for the mean expression level of gene *g* based on the observed counts 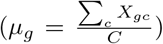 includes biological as well as technical effects. However, we are interested in inferring the mean expression level and variance that is primarily reflective of the underlying biology. This can be accomplished by relying on the size factors (equation. (7)) to adjust the observed counts of the respective cells and then obtaining the estimates for mean and variance based on the scaled counts. Thus,

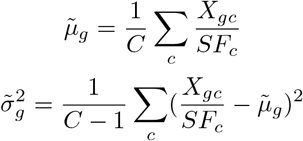

Using the estimates for mean and variance for gene *g*, we get the following estimates for mean and variance of gene *g* in each cell *c*,

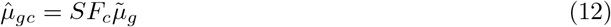

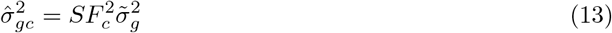

With these estimated means 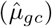 and variances 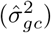 that account for the systematic biases, from equations. (10) and (11) we get,

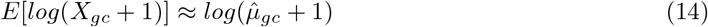

and

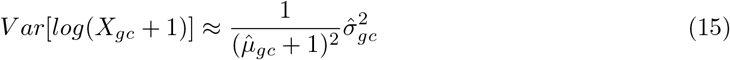

Based on these first-order approximations for means and variances under the *log*(*X* +1) transformation, we now define our *z*-scores based normalization,

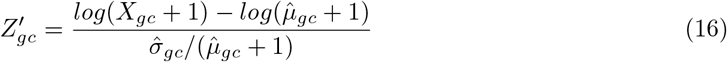

This *log*-stabilized *z*-score transformation is the default normalization method in our R package called `Piccolo` (see Appendix for a discussion on other variance stabilization approaches implemented in `Piccolo`). Hereafter, we refer to it as the `Piccolo` normalization.

#### Piccolo normalization reduces the impact of sampling depth differences between cells while simultaneously ensuring variance stabilization

As stated earlier, the objective of normalization is to reduce or eliminate the systematic biases in counts between cells that are not reflective of actual biological differences. Since technical control data do not have any biological source of variation, differences in sampling depths are expected to be the major source of variation between the cells (droplets). With PCA, this would translate to sampling depth showing up as a major contributor in one of the first few principal components (PCs).

In Fig. 3A, we show the scatter plots of cells based on their coordinates along PC1 and PC2. The colors of the dots reflect the size factors of the respective cells, with brighter shades (yellow) indicating larger size factors. Recall that larger size factors correspond to cells with larger sampling depths across the stable genes (see equation. (7)). In order to evaluate whether size factors correlate with the first few PCs, we calculated the canonical correlation coefficient (*ρ*) [31] between the size factors and the top 5 PCs; *ρ* close to 1 would indicate strong correlation between the size factors and one of the top 5 PCs. For `Svensson 1` raw counts (left panel Fig. 3A), we can clearly observe a color gradient along PC1, with cells with larger size factors lying predominantly on the left and cells with smaller size factors predominantly on the right. Thus for raw counts, sampling depth differences indeed show up as a major source of variation between the cells. The value of the canonical correlation coefficient - *ρ* = 0.97 - supports this further.

**Figure 3.**
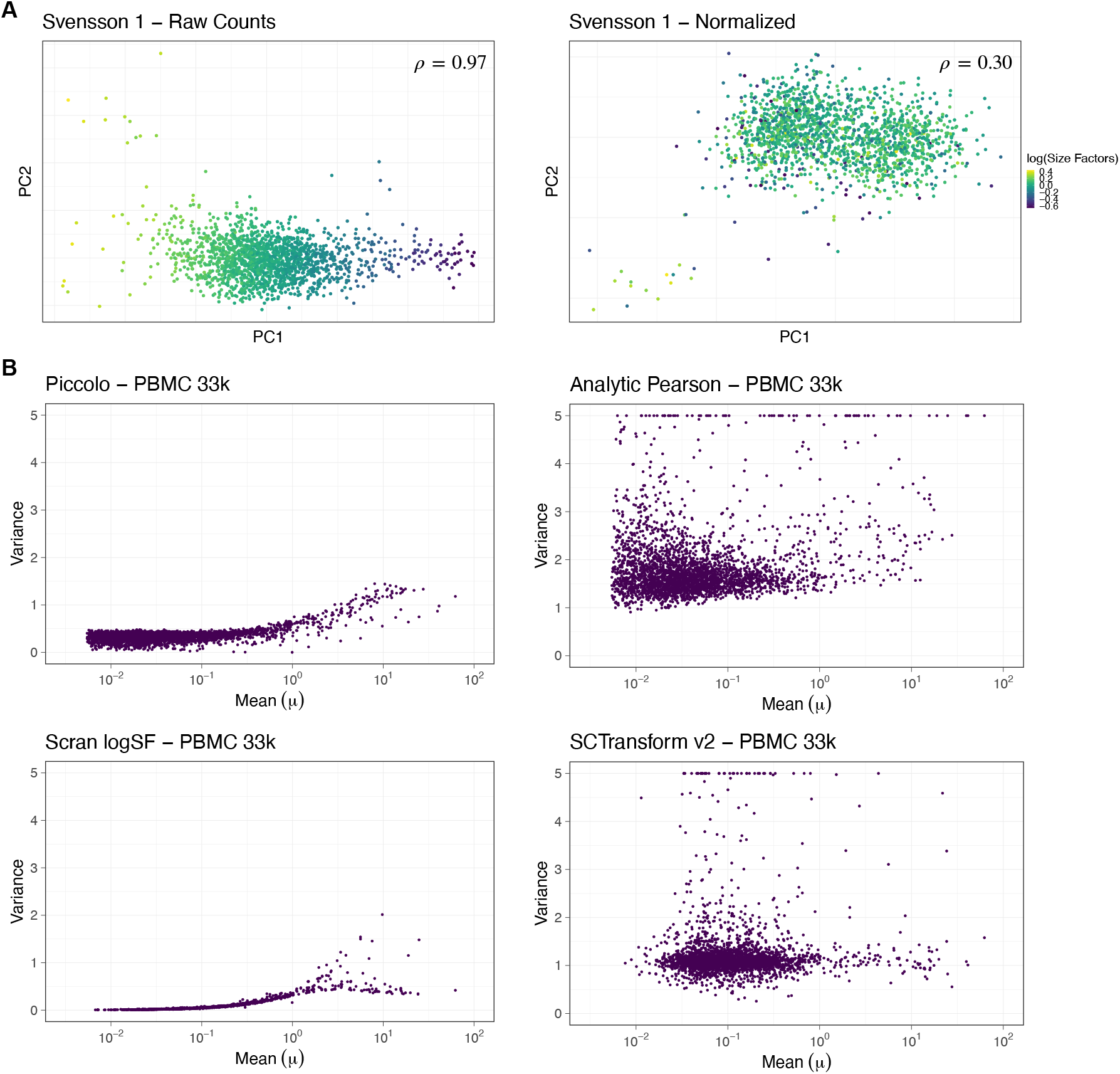
Piccolo normalization reduces sampling depth differences between cells and also ensures effective variance stabilization. **A**. 2-dimensional (2D) scatter plots based on the first 2 PCs of the Svensson 1 technical control data set. Each dot is a cell and is colored according to the size factors; brighter shades (yellow) correspond to larger size factors and darker shades (deep blue) correspond to smaller size factors. The left panel shows the 2D PC scatter plot for the raw counts, while the right panel shows the 2D PC scatter plot for the *z*-scores (residuals) obtained from Piccolo normalization. The coefficient (*ρ*) in the top-right corner of the panels shows the canonical correlation coefficient between the size factors and the top 5 PCs. Smaller values of *ρ* indicate that the impact of the sampling depth differences has been reduced more effectively by the normalization. Piccolo normalization reduces the impact of sampling depth on the overall variation as evaluated through PCA. **B**. Variance vs mean (*μ*) linear-log scatter plots for the top 3000 HVGs of the PBMC 33k data set after applying respective normalizations - Piccolo (top-left panel), Analytic Pearson residuals (top-right), Scran logSF (bottom-left), and SCTransform v2 (bottom-right). The *y*-axis scale was limited to a maximum value of 5 to aid visual comparison; genes with variance greater than 5 were clipped to have the maximum value of 5. The residuals obtained using Piccolo exhibit variances that do not vary much beyond 1 unlike the the residuals obtained with Analytic Pearson and SCTransform v2.

Next, we applied the `Piccolo` normalization to `Svensson 1` (right panel in Fig. 3A) and confirmed that the sampling depth differences are no longer identified as a major source of variation by PCA. In fact, not only do we not observe a color gradient along PC1 or PC2, but even the canonical correlation coefficient between the size factors and the top 5 PCs is significantly reduced to *ρ* = 0.3 which suggests a weak correlation. Similar observations were made for another technical control data set [32] (see Fig. S17). These results demonstrate that the `Piccolo` normalization is able to reduce the impact of systematic differences in sampling depths.

To examine the effectiveness of variance stabilization, we looked at the variances of the residuals after applying normalization to the raw counts. We compared `Piccolo` with two other residuals-based normalization methods - analytic Pearson [21](`Analytic Pearson`), and the regularized NB regression approach in scTransform [20, 29] (`SCTransform v2`). For reference, we also looked at the simple `logSF` based normalization approach implemented in Scran [12] (`Scran logSF`). Note, Scran relies on pooling of cells to arrive at estimates for the cell-specific size factors that are then used for the `logSF` normalization.

For the `PBMC 33k` data set, we first used our feature selection method to shortlist the top 3000 HVGs, and then applied the `Piccolo` normalization to compute the residuals corresponding to the raw counts of these HVGs. The `Analytic Pearson` residuals were also computed for these top 3000 HVGs identified with our feature selection method. This enables a direct comparison between the two methods since the residuals were calculated using the same set of features. In contrast, for `SCTransform v2` and `Scran logSF`, the top 3000 HVGs were shortlisted using their own respective approaches. For each of the normalization methods, we then calculated the variances of the residuals. These variances are shown in the variance-mean linear-log plots in Fig. 3B. It is apparent that the residuals obtained from `Piccolo` (Fig. 3B, top-left panel) exhibit much lesser scatter compared to the other two residuals-based approaches (Fig. 3B, top-right panel and bottom-right panel). In Fig. 3B bottom-left panel, we also show the variance of the *log*-transformed normalized values obtained with `Scran logSF` for reference. We note that the log-transformed values also exhibit much lesser deviation from 1 compared to the raw counts based residuals methods. Note, given the heteroskedastic nature of our counts data the increase in the variances of the transformed values (obtained from log-transformation based normalization or our variance stabilized residuals based normalization) as the mean expression levels increase is not eliminated. However, compared to the raw counts this dependence is reduced for the transformed values which plays an important role when we employ PCA downstream.

### Piccolo feature selection and normalization lead to substantial improvements in cell clustering

#### Residuals-based normalization which includes a variance stabilization transformation preserves cell-cell similarities between cells that share cell-type identities

Normalization is typically followed by dimensionality reduction and unsupervised clustering to identify groups of cells with similar expression profiles. Depending on the biological system, the groups of cells may correspond to distinct cell-types, or states. The identification of such groups is a pivotal step since it informs crucial downstream analyses such as differential expression and marker genes identification. The most popular scRNA-seq workflows (for example, Seurat [15–18] and Scanpy [19]) employ PCA to perform dimensionality reduction. Based on the PCs, *k*-nearest neighbour (*k*-NN) graphs are generated (with cells as nodes) in which communities of cells that are most similar to each other are detected using graph-partitioning algorithms such as Leiden [33] and Louvain [34].

To investigate how well our normalization method preserves cell-cell similarities between cells that share cell-type identities, we began by examining a truth-known data set (data set in which the cell-type identities of the cells are already known) prepared by Duo et al. [35] using cells purified with cell-type specific isolation kits by Zheng et al. [36]. Briefly, they prepared the data set by shortlisting purified cells belonging to 8 PBMC cell-types - B-cells, CD14 monocytes, CD56 NK cells, CD4 T-helper cells, memory T-cells, naive T-cells, naive cytotoxic T-cells, and regulatory T-cells - such that there were roughly equal numbers of cells corresponding to each cell-type in the final data set (between 400-600 cells per cell-type). We refer to this data set as `Zheng Mix 8eq`. Since `Zheng Mix 8eq` consists of a mix of well-separated cell-types (for instance, B-cells vs T-cells) and similar cell-types (different types of T-cells), it provides a simple yet reasonably challenging scenario for evaluating how well cells belonging to the different cell-types can be distinguished after normalizing the counts with the respective normalization methods.

We used `Piccolo` to shortlist the top 3000 HVGs and applied our normalization to obtain the residuals for those HVGs. For `Analytic Pearson`, the residuals were computed for the top 3000 HVGs shortlisted using our feature selection method, while for `Scran logSF` and `SCTransform v2` the transformed counts and residuals were respectively computed for the top 3000 HVGs shortlisted using their own methods. Subsequently, we performed PCA and shortlisted the first 50 PCs. We then used a simple *k*-NN based classification approach based on the PCs to predict cell-type labels for each cell by relying on the known cell-type labels (see Methods). Finally, we evaluated the extent of the agreement between the predicted labels and the known labels by calculating the following clustering metrics: the Macro F1 score, the adjusted Rand index (ARI), and the adjusted mutual information (AMI).

In Fig. 4A, we show the Uniform Manifold Approximation and Projection (UMAP) [37] plots for `Zheng Mix 8eq` with `Piccolo` normalization (top-left panel), `Analytic Pearson` (top-right panel), `Scran logSF` (bottom-left panel), and `SCTransform v2` (bottom-right panel). In the plots, each dot represents a cell and is colored according to the known cell-type labels (legend provided at the bottom of the 4 panels). Qualitatively, it is apparent from the UMAP plots that the B-cells, CD56 NK cells, and CD14 monocytes can easily be distinguished compared to the rest. As expected, it’s more difficult to distinguish between the different kinds of T-cells, with memory and naive cytotoxic T-cells being the only ones that are comparatively easier to distinguish, particularly with the residuals-based approaches. The values of ARI, AMI, and Macro F1 quantifying the extent of the agreement between the predicted and the known cell labels are listed in the bottom-right corner of the panels for the respective normalization methods. While there isn’t a significant difference in the metrics between the 4 normalization methods, we do observe the highest values with `Piccolo`.

**Figure 4.**
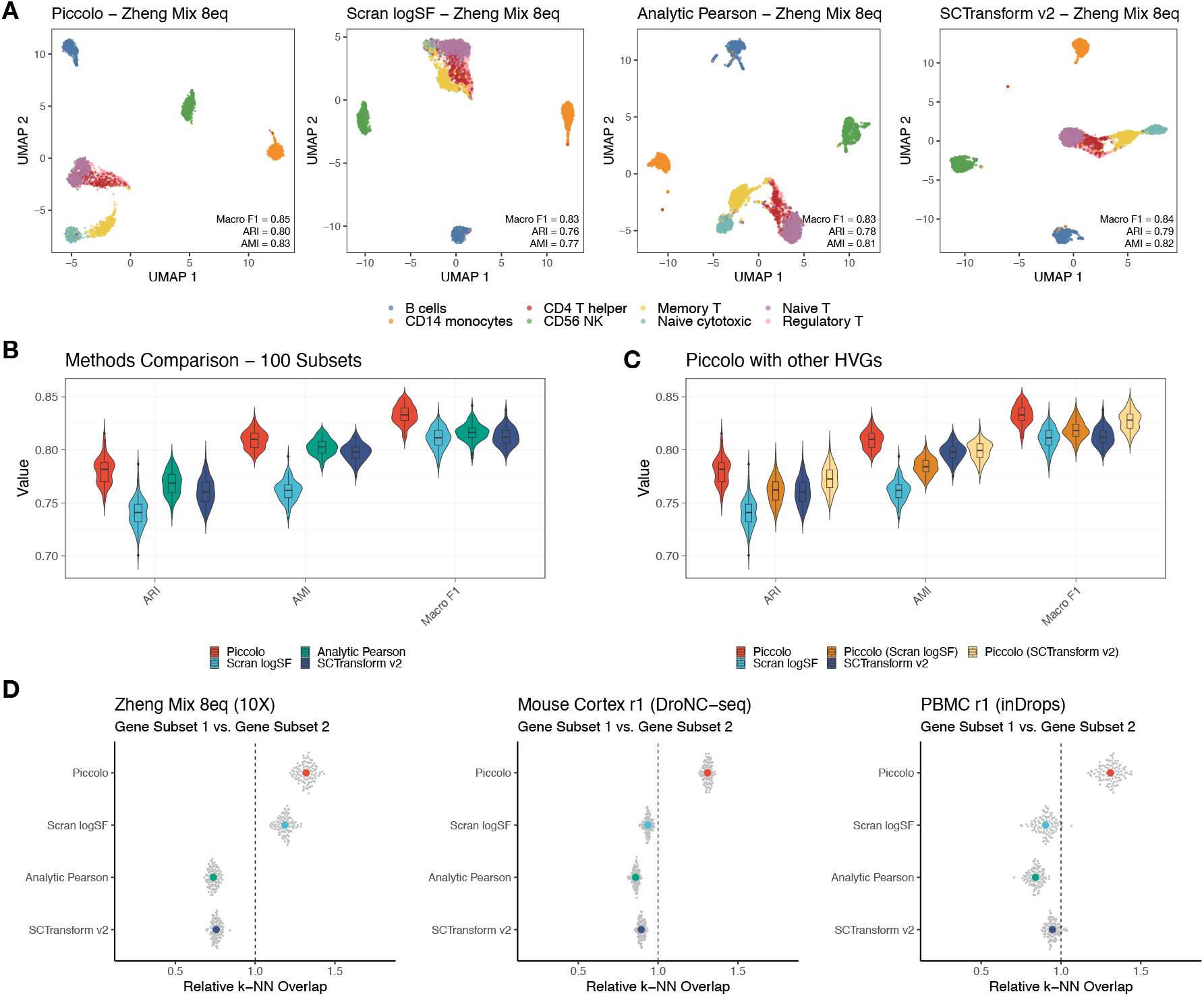
Piccolo normalization preserves cell-cell similarities between cells sharing cell-type identities. **A**. 2D UMAP plots after applying respective normalizations for the top 3000 HVGs of the Zheng Mix 8eq data set (consists of 3994 cells with roughly equal numbers of cells belonging to 8 distinct PBMC cell-types). Dots represent cells and are colored using the known cell-type labels (legend at the bottom of the panel). Clustering metrics - ARI, AMI, Macro F1 - based on comparisons between predicted cell labels (obtained using a kNN-based classification approach) and known cell labels are listed in the bottom-right corner in each panel. **B**. Violin-box plots of the clustering metrics obtained for 100 subsets of the Zheng Mix 8eq data set. The colors correspond to the respective normalization methods used. For all the 100 subsets, the highest values of the metrics was observed with Piccolo (red) (see Supplementary Table 1). **C**. Violin-box plots comparing the clustering metrics obtained for 100 subsets of the Zheng Mix 8eq data set with Piccolo normalization used with HVGs shortlisted obtained from other methods; Piccolo (Scran LogSF) and Piccolo (SCTransform v2) refer to the methods where the top 3000 HVGs were obtained using Scran logSF and SCTransform v2 respectively, and then the normalization was performed on those HVGs using Piccolo. Piccolo yielded higher values of the clustering metrics despite relying on HVGs shortlisted by other methods (see Supplementary Table 2). **D**. Comparisons of overlap between the *k*-NNs inferred separately on two halves of data obtained by randomly splitting genes evenly (done 100 times to create 100 pairs of split datasets). Relative *k*-NN overlap was calculated by dividing the mean overlap per data set by its average across all normalization methods. Colored dots indicate averages across the 100 splits (small grey dots) per normalization method - their colors are consistent with the colors in panel B. This panel is similar to Fig. 2a in [22] and highlights that Piccolo (red) surpasses the other methods in maintaining *k*-NN consistency.

However, these observations are not sufficient to argue for or against any of the methods. For a more robust comparison between the normalization methods, we created 100 subsets of `Zheng Mix 8eq` by randomly picking 50% of the cells in the original data set 100 times. For each of the 100 subsets, we used the same approach as discussed above for the 4 normalization methods and computed the respective ARI, AMI, and Macro F1 scores for the predicted cell labels based on the *k*NN-based classification approach. In Fig. 4B, we show the violin-box plots for the ARI, AMI, and Macro F1 scores for the 100 subsets. The colors correspond to the respective normalization methods used (red corresponds to `Piccolo`). We can clearly see that for all 3 clustering metrics, the highest values were consistently obtained with `Piccolo` normalization, reflecting that the best agreement between predicted and known labels is achieved using our feature selection and normalization method. For each clustering metric, we used paired Wilcoxon tests to quantify whether the differences between the values of the metric obtained with `Piccolo` normalization and other normalization methods were statistically significant (see Methods). For all 3 metrics, the values obtained with `Piccolo` normalization were found to be consistently higher than those obtained with other methods (all paired Wilcoxon test *p*-values were found to be highly significant - *p <* 1*E*− 11, see Supplementary Table 2).

For our analyses on `Zheng Mix 8eq` so far, while `Piccolo` and `Analytic Pearson` normalization were applied on the same set of HVGs, the sets of HVGs for `Scran logSF` and `SCTransform v2` were different. Thus, some of the differences in the results are attributable to the differences in the sets of HVGs. To compare just the normalization methods, we used the top 3000 HVGs shortlisted by `Scran logSF` and `SCTransform v2` respectively, and computed the residuals using `Piccolo` normalization for these respective HVGs. `Piccolo (Scran logSF)` denotes the method wherein the HVGs were obtained from `Scran logSF` and then the `Piccolo` normalization was performed for those HVGs. Similarly, `Piccolo (SCTransform v2)` denotes the method wherein the HVGs were obtained from `SCTransform v2` and then the `Piccolo` normalization was performed for those HVGs. By applying these methods, we arrived at clustering metrics for the 100 subsets which are shown using the violin-box plots in Fig. 4C. The colors correspond to the feature selection and normalization methods used. We used paired Wilcoxon tests to compare the values of the metrics obtained with the normalization method that was used to shortlist the HVGs, with values of the metrics obtained with `Piccolo` normalization using those HVGs. For ARI and Macro F1, the values obtained with `Piccolo` normalization (`Piccolo (Scran logSF)` and `Piccolo (SCTransform v2)`) were consistently higher than the values obtained with the respective normalization methods with which the HVGs were shortlisted (paired Wilcoxon test *p <* 1*E* − 09). For AMI, while the values with `Piccolo (Scran logSF)` were consistently higher than with `Scran logSF` (paired Wilcoxon test *p <* 1*E*− 17), the values obtained with `Piccolo (SCTransform v2)` were not as significantly high compared to the ones obtained with `SCTransform v2` (paired Wilcoxon test *p <* 0.06) (see Supplementary Table 3). Overall, it is clear that even with other HVGs, `Piccolo` normalization is better at preserving cell-cell similarities between cells of the same cell-type compared to the other normalization methods. Moreover, from Fig. 4C what is most striking is that the values of the clustering metrics obtained using our feature selection and normalization are the highest overall (red violin-box plot corresponds to `Piccolo`). This clearly suggests that not only is the performance of `Piccolo` normalization consistently better than the other normalization approaches (assessed with the aid of the clustering metrics), but even our feature selection method is effective at shortlisting HVGs that better inform the differences between cells with distinct cell-type identities.

#### Piccolo maintains consistency of *k*-nearest neighbors - a necessity for ensuring robustness of downstream analyses involving nearest-neighbor graphs

In a recent article comparing different transformations for scRNA-seq data, Ahlmann-Eltze and Huber highlighted the *k*-nearest neighbor (*k*-NN) graph as a fundamental data structure which is used to infer cell-types, states, trajectories etc [22]; to remind, the *k*-NN graph in scRNA-seq analyses is obtained by relying on a lower-dimensional representation of the cells using PCA, and subsequently shortlisting the *k*-NNs of each cell based on Euclidean distances in the PC space (typically *k* is 10). They pointed out that the consistency of the *k*-NNs is a necessary (albeit not sufficient) condition for the robustness of downstream analyses that rely on *k*-NN graphs. They evaluated *k*-NN consistency by evenly splitting each data set into two halves based on the genes, such that the two resultant subsets contained mutually exclusive sets of genes (they referred to them as gene subset 1 and gene subset 2, see Fig. 2a in [22]). They then applied the respective transformation approaches to the gene subset 1 and gene subset 2 datasets separately and obtained *k*-NNs for each cell corresponding to each of the subsets. Using these 2 sets of *k*-NN cells for each cell, they performed a pairwise comparison to determine the extent of overlap between them and thereby assess consistency.

We performed a similar analysis for the `Zheng Mix 8eq` (10X), `Mouse Cortex r1` (DroNC-seq), and `PBMC r1` (inDrops) datasets (see Methods). While Ahlmann-Eltze and Huber only considered 10X derived UMI counts datasets, we included datasets obtained from other droplet-based high-throughput technologies to showcase the performance of Piccolo in preserving *k*-NN consistency. We performed the gene-based splits for each data set 100 times. *k*-NN overlaps were obtained per cell and then averaged across all the cells to arrive at one mean estimate per iteration. Relative *k*-NN overlap values were calculated by dividing these mean NN overlap values by their average across all iterations for all 4 normalization methods. Fig. 4D shows the resultant values of the relative *k*-NN overlaps for each normalization method for the 100 splits (small grey dots). The large colored dots indicate the averages across the 100 splits per normalization method; colors of the dots were kept consistent with the colors used in panel B for each of the methods. Unlike Fig. 2a in [22] where the authors aggregated the relative *k*-NN overlap values across the datasets, we show the relative *k*-NN overlaps for each data set separately to highlight that Piccolo (red in Fig. 4D) easily surpasses the other methods in fulfilling the necessary condition of *k*-NN consistency despite basic differences in these droplet-based high-throughput technologies (all paired Wilcoxon test *p*-values between Piccolo and other methods were less than 2.2*E* − 10). Based on these results, we conclude that our proposed normalization method ensures the consistency of *k*-NNs and will therefore enable more robust inferences to be drawn from downstream analyses that rely on *k*-NN graphs.

#### Piccolo enables identification of groups containing few cells as well as groups with cells that express fewer differentially expressed genes

To further investigate the performance of `Piccolo`, we utilized Splat [38], a simulation framework that relies on the gamma-Poisson distribution to simulate counts based on estimation of parameters for real UMI counts datasets. We generated simulated counts using the `NIH/3T3` data set for the following two scenarios:

- `Simulated Counts Scenario 1`: 6 groups with different number of cells per group, while keeping the probability that any given gene is picked to be differentially expressed the same for each group. The objective behind this simulation was to examine whether the groups with the fewest cells can be reliably identified after applying the different normalization methods.
- `Simulated Counts Scenario 2`: 6 groups with the same number of cells per group, but with different probability for any given gene to be picked to be differentially expressed within each group. The objective behind this simulation was to examine how well we can distinguish between cells belonging to distinct groups, especially the ones that have fewer differentially expressed genes.

We will discuss each scenario in turn now. For `Simulated Counts Scenario 1`, while the sizes of the groups were varied such that Group 6 and Group 5 had the fewest cells, the probability that any given gene is picked to be differentially expressed was kept fixed at the same value for all groups (see Methods). We adopted the same procedure as described in the previous subsection to prepare the UMAP plots for each of the 4 normalization methods - `Piccolo`, `Analytic Pearson`, `Scran logSF`, and `SCTransform v2` respectively (Fig. 5A). The cells were colored according to the group to which they belong. The legend for the group identities is provided at the bottom of Fig. 5A. Using the *k*NN-based classification approach, we predicted the group labels for every cell by relying on the known group labels. The clustering metrics quantifying the extent of the agreement between the predicted and the known group labels are listed in the bottom-right corner of the panels for the respective normalization methods. While the values of the clustering metrics clearly indicate the vast improvement enabled by `Piccolo` (highest values for all 3 metrics), even a qualitative inspection of Fig. 5A suggests that `Piccolo` performs quite favorably compared to the other normalization methods.

**Figure 5.**
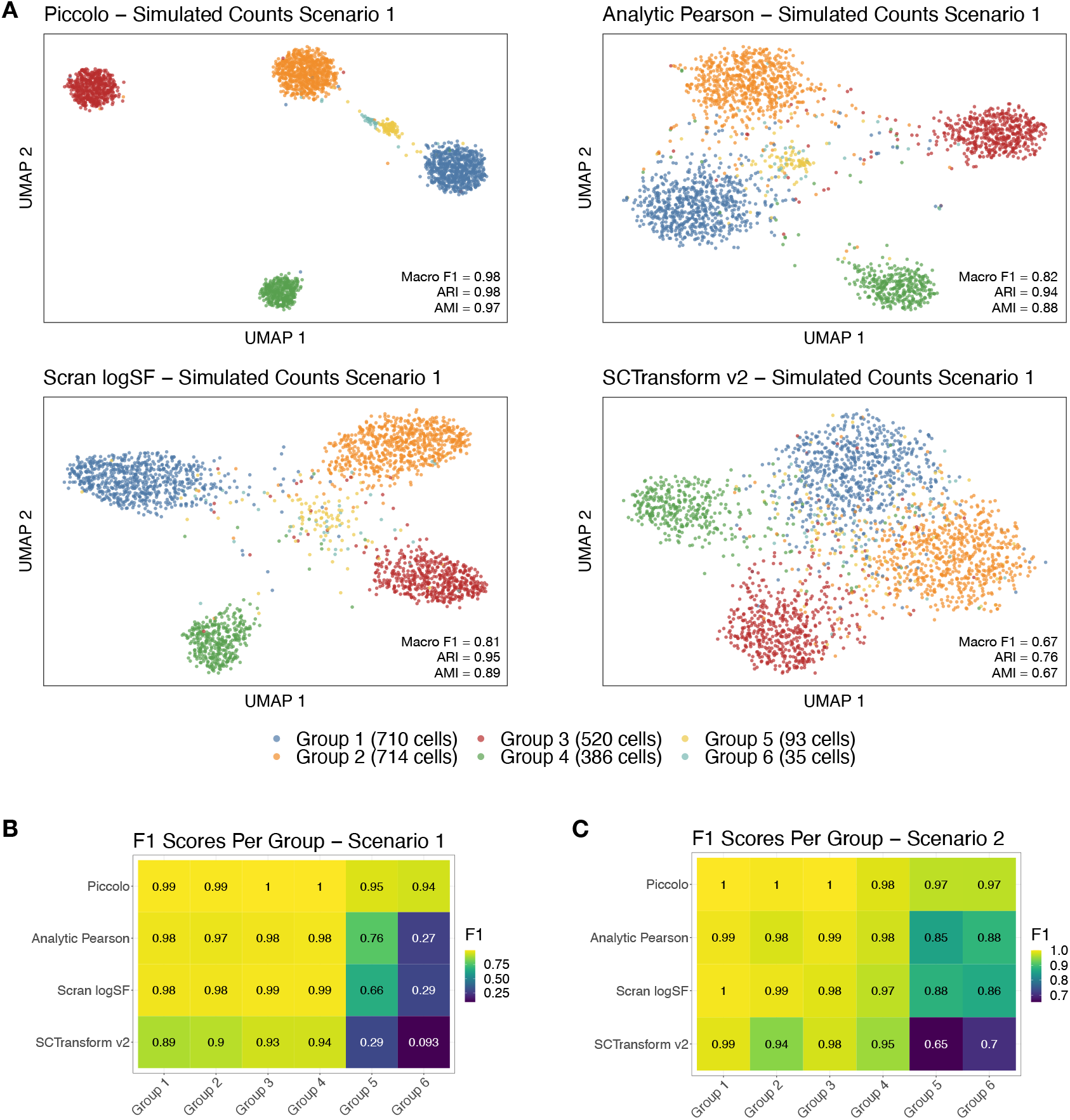
Piccolo enables identification of clusters even for groups containing small numbers of cells. **A.** 2D UMAP plots after using respective normalization methods for the top 3000 HVGs for the Simulated Counts Scenario 1 data set. The simulated counts were generated by Splat [38] using the NIH/3T3 data set. Dots represent cells and are colored using the known cell-type labels (legend at the bottom of the panel; actual numbers of cells belonging to each group are specified in parentheses). Clustering metrics based on comparisons between predicted cell labels and known cell labels are listed in the bottom-right corner in each panel. Cells belonging to the 6 groups can be more readily distinguished using Piccolo. **B.** Heatmap showing the F1 scores per group for each of the normalization methods applied to Simulated Counts Scenario 1. The tiles of the heatmap are colored according to the F1 values with larger F1 scores corresponding to brighter shades (yellow), and lower F1 scores corresponding to darker shades (deep blue). Piccolo has the highest F1 scores for all 6 groups. **C.** Heatmap of F1 scores per group for the Simulated Counts Scenario 2 data set (also see Fig. S13). Piccolo returns the highest F1 scores for all 6 groups. Note, the colors are scaled differently in panels B and C to bring out the differences between the F1 scores in the respective simulation scenarios.

The difference is especially stark for the groups containing the fewest cells (Group 5 - 93 cells, and Group 6 - 35 cells). With the aid of our feature selection and normalization (top-left panel in Fig. 5A), we observe in the resultant 2D UMAP embeddings that cells belonging to the respective groups form clusters which can be easily distinguished from clusters corresponding to other groups. For the other normalization methods, distinct clusters corresponding to the smallest groups are not as evident. In Fig. 5B, we show the F1 scores per group based on the concordance between the predicted and the known labels for each of the normalization methods respectively. The tiles of the grid have been colored based on the values of the F1 scores, with brighter shades (yellow) indicating larger values (greater agreement between the predicted and the known group labels), and darker shades (dark blue) indicating smaller values (less agreement between the predicted and the known group labels). The difference between `Piccolo` and other methods is quite stark, with `Piccolo` clearly better at preserving cell-cell similarities between cells sharing group identities. Note, our feature selection also plays a key role by shortlisting variable genes that enable the identification of distinct clusters that correspond to the distinct groups.

For `Simulated Counts Scenario 2`, we kept the number of cells in each group roughly the same and reduced the number of DE genes corresponding to each group such that Group 6 and Group 5 cells had the fewest DE genes. We used the same approach as the one described above. The resultant UMAPs are shown in Fig. S13. Once again, from just the qualitative differences in the nature of clustering of the cells in the respective UMAPs, it is clear that `Piccolo` performs quite favorably compared to the other normalization methods, especially when it comes to distinguishing between the groups with the fewest differentially expressed genes (Group 5 and Group 6). The clustering metrics reflect these differences in the overall quality of the clustering as well, with `Piccolo` exhibiting the highest values for all 3 metrics. In Fig. 5C, we show the F1 scores per group based on the predicted labels for each of the normalization methods respectively. `Piccolo` clearly does better at preserving similarities as well as distinctions between cells compared to the other normalization methods. In particular, cells belonging to groups with fewer differentially expressed genes can still be reliably distinguished with the help of `Piccolo`. We replicated the above simulation scenarios with another cell line data set (`HEK293T` [10]) with similar results that are summarized in Fig. S14 and Fig. S15.

In summary, we utilized a robust and popular simulation framework and derived single-cell counts based on 2 independent datasets for two distinct scenarios. The simulation scenarios were deliberately kept simple in order to facilitate a more straightforward comparison and interpretation. Despite the simplicity, we observed significant improvements in the clustering outcomes with the help of `Piccolo`. When viewed from the perspective of the identification of rare cell types (similar to Scenario 1), or differentiating between cell states (similar to Scenario 2), these results suggest that `Piccolo` feature selection and normalization will enable more robust scRNA-seq downstream analyses.

## Discussion

We began our investigation into the nature of UMI counts by examining the mean-variance relationships of the observed counts for the genes and showed that for genes with small counts the variance of the counts can be approximated quite well by the quasi-Poisson variance. We pointed out that this quasi-Poisson nature of the variance simply reflects the fact that the counts for the respective genes are small in most cells. We followed this by examining and questioning the assumption underlying typical scRNA-seq workflows. In a typical scRNA-seq workflow, feature selection is preceded by normalization. Implicit in this sequence of steps is the assumption that features which exhibit high biological variability can be identified only after taking into account the systematic technical biases. We pointed out that this assumption reflects a confusion between the distinct objectives of identification of the differentially expressed genes, and the HVGs. While it is imperative that the counts be normalized to account for the systematic biases prior to a differential expression analysis, we showed that in fact it is possible to identify HVGs based on just the observed counts. We proposed a simple approach for feature selection that relies on quasi-Poisson dispersion coefficients estimated from the observed counts using a regression-based method. A key advantage with assessing the overall variability of counts for each gene prior to normalization is the ability to identify genes whose counts do not vary significantly across the cells. We refer to these genes as *stable genes*. We posited that the variability of counts for such stable genes is primarily reflective of the systematic biases (such as sampling depth differences) and can more reliably inform the estimation of size factors for each cell. Based on these observations, we propose a revision of the scRNA-seq workflow in which feature selection precedes, and in fact informs normalization (see Fig. 6).

**Figure 6.**
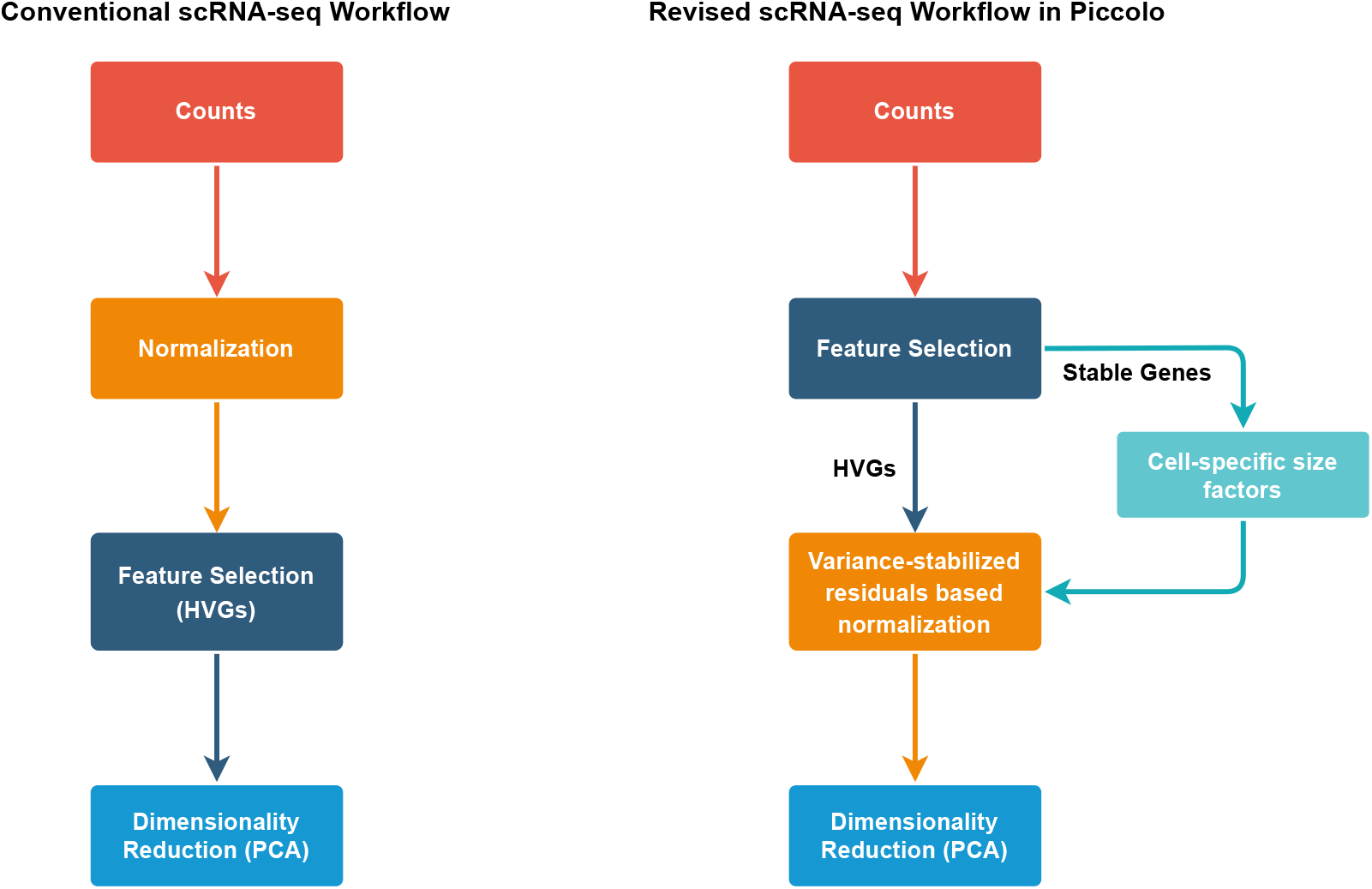
The revised scRNA-seq workflow implemented in Piccolo. We contrast the conventional scRNA-seq workflow (left) with the revised scRNA-seq workflow (right) proposed by us and implemented in Piccolo. The starting point for all analyses are the counts matrices. In the conventional workflow, the counts are first normalized and variance stabilized followed by feature selection to identify HVGs. Dimensionality reduction is then performed on the normalized counts. In contrast, in the revised workflow proposed by us, we first perform feature selection to identify both the HVGs as well as stable genes. We rely on the stable genes to estimate cell-specific size factors for performing normalization. During normalization, we also ensure variance stabilization which in turn leads to significant improvement while performing dimensionality reduction with PCA.

Before proceeding to discuss the salient aspects of our revised workflow and the normalization method proposed in this paper, it will be helpful to summarize and highlight some key aspects of the existing methods. In a recent article comparing different transformations for scRNA-seq data, Ahlmann-Eltze and Huber showed that the simple `logSF` normalization outperformed the residuals-based normalization methods particularly in ensuring consistency of the *k*-nearest neighbors [22]; consistency was evaluated by evenly splitting the data into two halves such that the two subsets contained mutually exclusive sets of genes, and inquiring whether the cells shared the same sets of neighbouring cells between the split subsets (nearest neighbour cells were identified based on Euclidean distances between the cells in the PC space). This consistency is primarily attributable to the variance stabilization ensured by the log-transformation. However, as pointed out earlier, a significant disadvantage of the `logSF` normalization is that it is not very effective at reducing the sampling depth differences between the cells (Fig. S8). In contrast, the residuals-based methods reduce sampling-depth differences between the cells much more effectively (Fig. S9). However, a drawback of the residuals-based approaches is that they are unable to effectively reduce and stabilize the variances, particularly for marker genes (see Appendix and [22]).

Keeping in mind the respective advantages and disadvantages of the `logSF` normalization and the residuals-based methods described above, we proposed a residuals-based (*z*-scores) based normalization method which includes a variance stabilization transformation (the default transformation function used in `Piccolo` is the *log*, see Appendix for descriptions of other options).

Given the observation that the majority of counts obtained from high-throughput technologies are small, we relied on first-order approximations for the means and variances of the transformed counts to estimate the residuals (see Appendix and Fig. S10). In addition, the residuals relied on means and variances that were adjusted using the size factors estimated with the help of stable genes. Thus, our conceptually straightforward approach simultaneously ensures variance stabilization and reduces the impact of the sampling depth differences thereby leading to concrete improvements in the downstream clustering performance.

We first applied our normalization method on technical control datasets (Fig. 3A and Fig. S11) and demonstrated that it does reduce the impact of sampling depth differences between the cells. Using the PBMC 33k data set (and other datasets, see Fig. S12), we further showed that our normalization method also ensures effective variance stabilization. We then applied `Piccolo` to a truth-known data set, and showed that it improves preservation of cell-cell similarities between cells that share cell-type identities. We also examined how well the consistency of *k*-NNs was ensured by `Piccolo`, and demonstrated that it consistently surpasses the other methods in satisfying this basic requirement for ensuring the robustness of downstream analyses that rely on *k*-NN graphs (see Fig. 4D). With the help of simulations, we were also able to show that `Piccolo` consistently enabled the identification of groups of cells that contain small number of cells, whereas other normalization methods failed to consistently and reliably ensure the same. In addition, we also showed that with `Piccolo` we can better distinguish between groups that express fewer differentially expressed genes. These results are especially relevant biologically when viewed from the perspective of the identification of rare cell-types, or distinguishing between cell states.

While our method offers significant improvements over the existing workflows, we now discuss some of its limitations. Beginning with feature selection, a fundamental assumption underlying our bin-based approach is that across all expression levels there are always some genes which are not biologically variable. This assumption is not motivated by biological observations and has been made to facilitate ease of computation. It is possible that we overlook some HVGs because of this assumption. In our current implementation, we mitigate this by keeping the level for the reference dispersion coefficient in each bin relatively low (10th quantile is the default). However, there is definite scope for further improvement of the feature selection process to ensure that the HVGs are more effectively identified. Another point of consideration tied to feature selection is the use of stable genes for estimating cell-specific size factors. Due to the sparsity of the data, it is possible that the counts across the stable genes for some cells are all zero resulting in the size factor estimates to be zero for those cells. In `Piccolo`, we address this by iteratively adding sets of genes from the bottom of the list of HVGs till none of the cells have a size factor estimate of zero. Despite this limitation, we still expect that these size factors will not be confounded by actual biological variation as much as when we estimate them using all the genes (since that includes the HVGs). With regard to our normalization method, we relied on the first-order approximations for both the mean and the variance under the variance stabilization transformation. These approximations will work well as long as the non-linear transformation function is approximately linear in the range of the observed counts. For small counts, this is indeed true, however for larger counts these approximations may lead to incorrect estimates. Given the nature of the droplet-based UMI counts data at present, our results suggest that the first-order approximations work quite well (see Appendix and Fig. S10). We point out here that for datasets that exhibit larger counts and less sparsity, the conventional approach of the log-based normalization can be expected to work reliably and is available as one of the options (called `logSF`) in the `Piccolo` R package. We also want to note that in this study we did not discuss and elaborate on differential expression (DE) analysis which forms a vital component of all scRNA-seq studies. This was done to focus attention on the conceptual clarifications and simplifications for the core steps of feature selection and normalization that shape all downstream analyses, including identification of DE genes. We remind here that if we are unable to consistently and reliably identify groups of cells that actually correspond to distinct cell-types or states, then the downstream DE analyses are unlikely to be as informative and helpful. In the worst cases, they may even be misleading. Briefly, we would like to mention here that after applying our normalization method to the observed counts, the distribution of the residuals are brought closer to the normal distribution, thus making it possible to employ the two-sample Student’s t-test with the null hypothesis that the means of the two samples are the same. Typically, in scRNA-seq analyses the Wilcoxon rank-sum test, which is a non-parametric alternative to the two-sample t-test, is the preferred choice.

Another point of consideration for single-cell workflows is the time taken to perform the normalization, as well as the memory (RAM) that is used. For the latter, since our residuals-based normalization transforms a sparse matrix to a dense matrix, the amount of memory that is used for the post-normalization matrix will increase significantly. However, since we can shortlist the HVGs before normalization, the transformed counts matrix will only be generated for the HVGs thus requiring lesser memory than what would be needed if normalization was performed for all genes. With regards to the computation time, since the transformation is based on a simple analytical relation, it only took between 30 seconds (for `Svensson 1`) to 3 minutes (for `PBMC 33k`) to transform the observed counts to the residuals for each of the datasets used in this paper (see Methods for system configuration).

In conclusion, the novel scRNA-seq workflow proposed in this study offers a paradigm shift that not only leads to conceptual simplifications but also leads to substantial improvements in the downstream analyses. We expect that the implementation of this workflow in `Piccolo` will facilitate more cogent and impactful inferences to be drawn from future single-cell gene expression studies.

## Methods

### Data preparation and preprocessing

The datasets used in the this study are listed in the table in the Datasets section, along with the links to the sources.

#### Cell and gene filtering

Primarily, the only cell filtering applied for all datasets was to ensure that all the cells had non-zero total counts. However for the evaluation of the effectiveness of our normalization method using PCA, we filtered cells from the `Svensson 1` data set that had total counts 3.5 median absolute deviation away from the median total count. This was done to reduce the impact of outliers on PCA.

For all the datasets, we excluded genes that had fewer than 0.5% cells with non-zero counts. This is the default gene filtering employed in `Piccolo`.

#### Selection of HVGs

For all the analyses, unless specified otherwise, we shortlisted the top 3000 HVGs. For `Piccolo` and `Analytic Pearson` residuals-based normalization, the identification of HVGs was done using our dispersion coefficient-based feature selection method. For `Scran logSF` and `SCTransform v2`, the top 3000 HVGs were shortlisted using their respective approaches that rely on post-normalization transformed counts/residuals to identify the genes with largest variances of the transformed counts/residuals.

#### Dimensionality reduction using PCA

After selecting the top 3000 HVGs and performing normalization on the counts of the HVGs, we used PCA for dimensionality reduction and shortlisted the top 50 PCs. Prior to using PCA, for the residuals-based methods (including `Piccolo`) we centered the residuals at 0, but did not scaled to unit variance. For the `logSF` normalization, we did not center or scale the transformed counts.

### Kendall’s and Spearman’s rank correlation tests

Kendall’s and Spearman’s rank correlation tests were used to evaluate whether there is a statistical dependence between the quasi-Poisson dispersion coefficient (*α*_*QP*_) and the mean expression levels (*μ*) for genes with *μ <* 0.1. Both tests evaluate how well the relationship between two variables can be described using a monotonic function. For both tests, the correlation coefficients - denoted as *τ* and *ρ* respectively - indicate a statistical dependence if the values are close to +1 or − 1, while values of *τ* or *ρ* closer to 0 indicate the absence of such a statistical dependence. For these tests, genes with non-zero counts in fewer than 2.5% of cells in the respective datasets were excluded.

### Benchmarking cell-type clustering using *k*-NN based classification

Our *k*-NN based approach to predict cell labels using known cell labels is based on a very simple premise. After we normalize the observed counts and perform PCA, the expectation is that in the PC space the cells that share cell-type (or group) identities are close to each other. Thus, if we examine the nearest neighbours of a given cell belonging to a given cell-type (or group), we expect to find that the nearest neighbours are predominantly cells belonging to the same cell-type (or group). Based on this simple expectation, we predict cell labels for each cell by considering its *k* (default *k* = 15) nearest neighbours and using the known cell-type (or group) labels to identify the cell-type (or group) that is most over-represented in the nearest neighbour set - the most over-represented cell-type is the predicted cell-type label for the given cell. Over-representation is assessed using the hypergeometric test. By testing for over-representation, we ensure that there is no bias against cell-types (or groups) that have fewer cells while predicting the cell-type identity for any given cell.

### Comparing clustering metrics obtained using the different normalization methods

For the 100 random subsets generated using the `Zheng Mix 8eq` data set, we applied the respective normalization methods and performed dimensionality reduction using PCA. We used the top 50 PCs for each of them, and with our kNN-based classification approach predicted cell-type labels for each cell that we then compared with the known cell labels using the following clustering metrics: Macro F1 (harmonic mean between precision and recall, averaged across the classes), adjusted Rand index (ARI), and adjusted mutual information (AMI). We compared the values of each of these metrics obtained with the different normalization methods by using the paired Wilcoxon rank-sum test. The null hypothesis is that there is no difference in the values of the metrics between the two groups being compared. The paired-test is essential in this context since the values of the metrics have to be compared pairwise for the same subset, and cannot be compared across different subsets.

The same 100 subsets were used to also evaluate how well `Piccolo` performs when we rely on HVGs shortlisted based on other normalization methods. Once again, we relied on the paired Wilcoxon rank-sum test to assess whether there are differences in clustering performance as evaluated through the clustering metrics: Macro F1, ARI, and AMI.

### Evaluating *k*-nearest neighbors consistency

For the `Zheng Mix 8eq` (10X), `Mouse Cortex` r1 (DroNC-seq) and `PBMC r1` (inDrops) datasets, we randomly split the genes into two even subsetted datasets - Gene Subset 1 and Gene Subset 2. We normalized the observed counts in each subset using the four methods discussed in the paper. We performed PCA on the normalized values to shortlist the top 50 PCs and identified the 10 nearest neighbors of each cell by relying on Euclidean distances between pairs of cells in the 50 dimensional PC space (dbscan [39, 40] was used to identify the NN cells). For each cell, we thus obtained two sets of 10 nearest neighbors from the respective subsets and quantified the extent of their overlap. We then took the mean of these per cell overlaps across all cells in the given data set to calculate the mean overlap for any given normalization method.

This procedure was repeated 100 times and the mean overlap values were recorded for each iteration. To obtain the relative *k*-NN overlap values, we took the ratio of each of the mean overlap value obtained from a given iteration for each normalization method with the average of all the mean overlap values across all iterations and all normalization methods. This yielded the relative *k*-NN overlap values shown in panel D in Figure. 4.

### Simulations

To generated simulated counts datasets, we used Splat [38], a simulation framework that relies on the gamma-Poisson distribution to simulate counts based on estimation of parameters for real UMI counts datasets. Splat allows for single-cell counts simulation wherein we can specify both the number of groups of cells, as well as the extent of differentially expressed genes in each group (this is specified in terms of the probability that a gene will get picked to be differentially expressed in each group). In order to generate the simulated counts, it requires a homogeneous cell population based on which the parameters for the simulation are estimated. Thus, we used 2 cell line datasets - `NIH/3T3` and `HEK293T` - to simulate the counts since the cells in these datasets constitute biologically homogeneous cell populations. Below we provide details of the parameters specified for each of the the 2 simulation scenarios described in the main text:

- `Simulated Counts Scenario 1`: For each reference data set (`NIH/3T3` and `HEK293T`), we simulated 6 groups containing cells in the following proportions: Group 1 - 0.30, Group 2 - 0.30, Group 3 - 0.20, Group 4 - 0.15, Group 5 - 0.04, and Group 6 - 0.01. The probability that any given gene was picked to be differentially expressed was kept fixed at the default value of 0.1 for all the groups (parameter `de.prob` in Splatter). Thus, Group 6 and Group 5 have the fewest cells.
- `Simulated Counts Scenario 2`: For each reference data set (`NIH/3T3` and `HEK293T`), we simulated 6 groups containing roughly the same numbers of cells. But this time, we varied the probability that a gene will be picked to be differentially expressed per group (`de.prob`) as follows: Group 1 - 0.25, Group 2 - 0.2, Group 3 - 0.15, Group 4 - 0.1, Group 5 - 0.05, Group 6 -0.025. Thus, Group 6 and Group 5 have the fewest differentially expressed genes.

### System configuration and software used

We used a Macbook Pro with the Apple M1 pro chip and 16GB RAM for all the analyses in this paper. `Piccolo` was developed using R (version 4.2.0), and all the analyses were also performed using R [41]. We used the following R packages in this study: cluster [42], data.table [43], dbscan [39, 40], ggplot2 [44], igraph [45], Matrix [46], matrixTests [47], RSpectra [48], Rtsne [49–51], umap [52], viridis [53]. In addition, the `Piccolo` package also includes a helper function made by Kamil Slowikowski called writeMMgz to help prepare .mtx.gz files.

## Datasets

| Name | Technology | Tissue | Type | Source | Reference |
| --- | --- | --- | --- | --- | --- |
| Svensson 1 | 10X Chromium v1 | Technical Control | Technical Control | Link | [10, 54] |
| Klein 2015 | inDrops | Technical Control | Technical Control | Link | [10, 32] |
| NIH/3T3 | 10X Chromium v3 | 3T3 | Cell Line | Link | [10] |
| HEK293T | 10X Chromium v3 | HEK | Cell Line | Link | [10] |
| PBMC 33k | 10X Chromium v1 | PBMC | Heterogeneous | Link | [20] |
| PBMC r1 | inDrops | PBMC | Heterogeneous | GSE132044 | [8] |
| Mouse Cortex | DroNC-seq | Cortex | Heterogeneous | GSE132044 | [8] |
| Mouse Lung | Drop-seq | Lung | Heterogeneous | GSE124872 | [25] |
| Zheng Mix 8eq | 10X Chromium v1 | PBMC | Heterogeneous | Link | [35, 55] |
| Haber 2017 | 10X Chromium v1 | Mouse Small Intestine Epithelium | Heterogeneous | GSE92332 | [56] |

## Data availability

All the datasets used in this study, including the simulated counts datasets, are publicly available at https://github.com/Amartya101/PiccoloPaperData

## Code availability

All the code used to perform the analysis and generate the figures in this paper are publicly available at https://github.com/Amartya101/PiccoloPaperData

## Availability of software

The development version of `Piccolo` is currently available as an R package through GitHub at https://github.com/Amartya101/Piccolo, and is licensed under the GNU GPLv3 license. In order to facilitate easier adoption with existing tools, the feature selection and normalization methods implemented in `Piccolo` can be used with Seurat (version 4) [18], the instructions are provided at https://github.com/Amartya101/Piccolo-With-Seurat

## Acknowledgments

We thank the members of the Khiabanian and Herranz labs for their encouragement and insightful feedback. Special thanks are due to Mona Arabzadeh and Vaidhyanathan Mahaganapathy for carefully reading the manuscript and suggesting improvements. This work was supported by grants from the National Institutes of Health (R01CA233662 and P30CA072720), the V Foundation (T2019-01), and the New Jersey Commission on Cancer Research (COCR23PRG006).

## Appendix Regression-based approach to estimate QP dispersion coefficients

The regression-based test proposed by Cameron and Trivedi [23, 24] relies on the idea that under the null hypothesis - counts, *x*, are Poisson distributed - the expected value of (*x* −*E*[*x*])^2^ −*x* is zero, while under the alternative hypothesis the expected value would be a scalar multiple of a function of *E*[*x*]. Since *E*[*x*] is unknown, we replace it by the estimate under the null hypothesis and estimate the scalar multiple using least squares regression.

The choice of the QP mean-variance relation corresponds to the alternative hypothesis in which the variance (*V ar*[*x*]) is a linear function of the mean (*E*[*x*]),

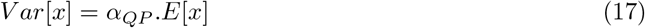

Given this specification of the alternative hypothesis, the *α*_*QP*_ are estimated by auxiliary ordinary least-squares regression using,

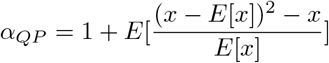

with *E*[*x*] = *μ* = Poisson estimate for the expected value of the counts.

### QP variance as a special case of NB variance and the manifestation of a non-decreasing relationship between *θ* and *μ*

It is instructive and useful to view QP variance as a special case of NB variance wherein *α*_*NB*_ exhibits dependence on *μ* through the following relationship

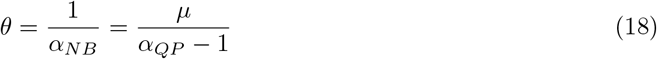

Based on the relation above, we expect that if the values of *α*_*QP*_ for genes with *μ* within a given range of mean expression levels do not exhibit a dependence on their *μ* (genes with larger *μ* do not necessarily have larger *α*_*QP*_), then we would observe a monotonic dependence between *θ* and *μ* (*θ* ∝ *μ*) for the genes with means within that range of mean expression levels.

Using equation. 18, we obtained estimates of *θ* for each gene using the corresponding values of *α*_*QP*_ and *μ*. We plotted the *θ* vs *μ* log-log plots for the genes using these values. For `Svensson 1` (left panel in Fig. S1 C), we observe a clear non-decreasing relationship between *θ* and *μ*, particularly for genes with low mean expression levels. In order to make the monotonic increase even more apparent, we plotted estimates of *θ* obtained by fixing *α*_*QP*_ to the value of *α*_*QP*_ for the gene with the highest point density estimate in the the *θ* vs *μ* log-log plot (the dashed red line). Furthermore, we plotted contour lines (white curves) based on the point densities as a visual aid to infer how the density of the points (genes) varies depending on *μ*. If we imagine a closed contour loop as an ellipse, the straight line through the center of the ellipse that joins the two points furthest from the center is called the major axis of the ellipse. We refer to analogous lines for the contours as their longer axes. Supposing no dependence of *θ* on *μ*, the longer axes of the contours would be parallel to the *x*−axis (zero slope). Instead, we observe that the dashed red line of the estimated *θ* with fixed value of *α*_*QP*_ lies along the same direction as the longer axes of the contours, particularly for genes located in the region with high point density (bright yellow region). For `PBMC 33k` (right panel in Fig. S1 C), once again, despite greater variability in *θ* due to the inherent biological variability, the monotonic increase in *θ* with *μ* particularly for genes with low mean expression levels is still very evident. Particularly for genes located in the region with high point density (bright yellow region), the dashed red line of the estimated *θ* with fixed *α*_*QP*_ closely follows the direction of the longer axes of the contours. The `NIH/3T3` data provides an insightful contrast (middle panel in Fig. 1C) - while the monotonic increase in *θ* with increase in *μ* is evident for genes with low mean expression levels, there is a clear discrepancy between the slope of the dashed red line (estimated *θ* with fixed value of *α*_*QP*_) and the slope of the longer axes of the contours, particularly for genes in the region with the highest point density (bright yellow region). A closer examination reveals that for this data set the mean expression level (*μ*) of genes in the region of highest point density lie between 0.1 and 1, which is an order of magnitude higher than what we observe for `Svensson 1` and `PBMC 33k` (*μ* lies between 0.01 and 0.1 for genes in their respective regions of highest density). This difference actually stems from the differences in their respective sequencing depths - while the median total UMIs per cell in `Svensson 1` and `PBMC 33k` are 2309 and 1891 respectively, it is 15560 in `NIH/3T3`.

We must mention here that scTransform [20] relies on this non-decreasing relationship between gene abundance (*μ*) and the inverse over-dispersion parameter (*θ*) to perform regularization. In a recent paper [29], Choudhary and Satija argued that when modeling scRNA-seq data using a Gamma-Poisson distribution the inverse over-dispersion parameter (*θ*) does vary as a function of the gene abundance (*μ*), but that the true nature of this relationship can be masked for genes with low molecular counts. Their justification for such a relationship primarily rests on observations made for bulk RNAseq studies. However, for scRNA-seq counts data there appears to be a simpler explanation not linked to any underlying biological cause, namely the Poisson-like variance of counts for genes with low mean expression levels. This QP variance for low abundance genes holds not just for biological datasets but also for negative control datasets thus suggesting that there is no biological source for this non-decreasing relationship.

### Hoes does the number of bins impact the feature selection process?

The choice of number of bins plays an important role in the feature selection process. Fewer bins would lead to more genes per bin and this would result in an underestimation of the value of the quantile, especially for the bin that contains genes with the largest mean expression levels since their *α*_*QP*_ vary the most with *μ*. We show this in Fig. S6 for the `PBMC 33k` data set. When we pick the number of bins to be 10 (top left panel in Fig. S6), we observe a noticeable increase in the values of *α*_*QP*_ −*α*_*QP* (*Reference*|*Bin*)_ with *μ* for genes with the highest mean expression levels. This will lead to a bias towards selection of high expression genes as HVGs. We can increase the number of bins to reduce this bias since this will ensure that genes with comparable mean expression levels are being grouped together (see bottom left panel for 1000 bins). However, if we keep on increasing the number of bins there will be very few genes per bin. Since we effectively assume that in every bin there is a stable gene we would end up concluding for a significant proportion of genes that these genes are stable even if they actually exhibit biological variability (note the decrease in values of *α*_*QP*_ − *α*_*QP* (*Reference*|*Bin*)_ for genes with high mean expression levels in the bottom right panel; FTH1 for instance goes from having a value above 40 when the number of bins was set to 1000 to a value below 40 with the number of bins set to 10000). In practice, setting the number of bins to the default value of 1000 typically leads to 10-15 genes per bin with the gene with the smallest or the second smallest value within the bin providing the *α*_*QP* (*Reference*|*Bin*)_ value for that bin.

### The standard estimate of cell-specific size factors assumes that the counts are Poisson distributed

There is an intimate link between the estimates of size factors given by equation. (6) and the estimates for expected means 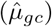 under the assumption that the counts are Poisson distributed. Assuming that each gene *g* contributes a proportion *p*_*g*_ of the total count *N*_*c*_ in cell *c*, the counts *X*_*gc*_ are modeled as,

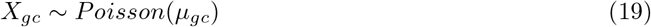

where *μ*_*gc*_ = *p*_*g*_*N*_*c*_. As shown by Townes et al. [27] and Lause et al. [21], the maximum likelihood estimates for *p*_*g*_ and *N*_*c*_ under the Poisson model are given by,

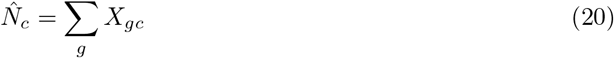

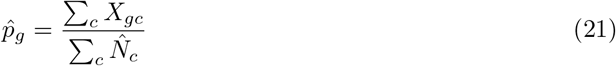

Based on these estimates, the maximum likelihood estimate for *μ*_*gc*_ is given by,

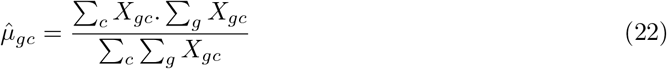

If we divide both the numerator and denominator in equation. (22) by the total number of cells (*C*), we get,

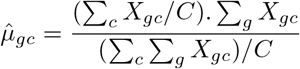

Since Σ_*c*_ *X*_*gc*_*/C* = *μ*_*g*_ = mean of the observed counts of gene *g*,

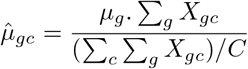

Using equation. (6), we can rewrite the above equation more simply as,

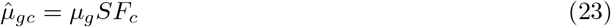

From this we can conclude that the simple estimates of the size factors given by equation. (6) should be more appropriately viewed as estimates under the approximation that the counts are Poisson distributed.

### Estimates for mean and variance under variance stabilization transformation

In order to compute *z*-scores for our data, we first need to apply a variance stabilization transformation to the observed counts (*X*_*gc*_) to bring their distribution closer to the normal distribution. The variance stabilization transformation can be performed using monotonic non-linear functions, *g*(*X*), such that the transformed counts are given by,

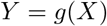

To compute the residuals, we need estimates for the means and variances of *Y* based on estimates for means and variances of *X*. We can arrive at approximations for both using a Taylor expansion around *X* = *μ*,

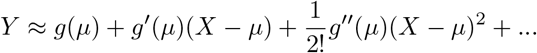

where *g*^′^(*μ*) and *g*^′′^(*μ*) are the first and second order derivatives of *g*(*X*) evaluated at *X* = *μ*. Considering the expansion up till the 1st order,

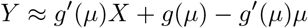

The expected value of *Y* can be approximated to,

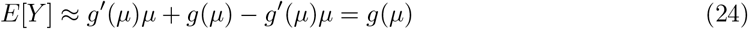

In addition, since

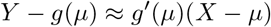

after squaring and taking expectation we get the following approximation for the variance of *Y*,

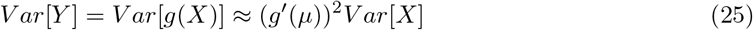

For *g*(*X*) = *log*(*X* + 1), the first order approximations of the mean and variance are then given by,

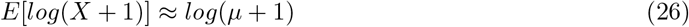

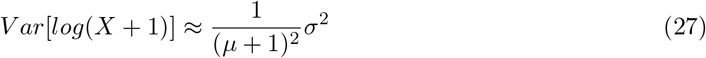

### First-order approximations for estimates of mean and variance under variance stabilization transformation are valid for majority of the genes

The first-order approximations for the mean and variance of the counts under the variance stabilization transformation in our residuals-based normalization are based on the assumption that the transformed values (raw counts transformed by the variance stabilizing transformation) vary linearly over the range of raw counts close to the mean expression level. We tested the validity of this assumption for each gene by fitting a straight line (using linear regression) through the *log*-transformed values corresponding to those raw counts that fall within one standard deviation away from the mean expression value. We relied on the resultant adjusted *R*^2^ (coefficient of determination) values to evaluate the validity of the assumption since values of adjusted *R*^2^ close to 1 indicate that the transformed values can be approximated to lie along a straight line over the corresponding range of raw counts.

We found that for all UMI counts datasets included in our study, more than 85% of the genes had adjusted *R*^2^ values greater than 0.8 (see Supplementary Table 3 and Fig. S10). Thus, for UMI counts data obtained using high-throughput protocols the first-order approximation utilized in our normalization method is valid, particularly for genes with small counts.

### Limitation of residuals without variance stabilization

We discuss the limitation of residuals computed for raw untransformed counts by considering the analytic Pearson residuals approach proposed by Lause et al. [21]. They proposed that the expected means 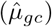 can be approximated with the estimates given by equation. (22). Further, they argued that the over-dispersion coefficient 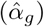 can be approximated with a fixed value of 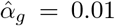 corresponding to the typical over-dispersion observed in technical control datasets. Their rationale for these choices is that the null model should correspond to the measurement process so that the residuals provide estimates for deviations compared to expectations under the measurement model. While conceptually sound, this approach compromises variance stabilization, especially for genes that are robustly expressed in only a subset of cells while showing negligible expression in the rest of the cells (such genes would be considered as *markers* of the specific cell sub-populations in which they are expressed).

We can illustrate the nature of the problem with the help of a simplified example. Suppose we have a data set consisting of 10000 cells with a gene (gene *A*) that is only expressed in 100 cells with identical counts of 100 in each of those cells; the rest of the cells have 0 counts. For simplicity, if we assume that all the cells have the same sequencing depth (*SF*_*c*_ = 1 ∀ *c*) then,

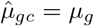

Based on this, equation. (8) simplifies to,

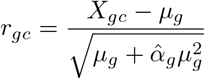

Given the distribution of counts for gene A stated above, *μ*_*A*_ = 1, and since we assume 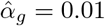, the residuals of gene A for the cells with counts of 100 are approximately 98.509, while the residuals for cells with 0 counts are -0.995. The overall variance of the residuals for gene *A* is approximately 98.03, thus exhibiting significant deviation from the null expectation of 1. The Pearson residuals method proposed by Hafemeister et al. [20] that allows for per gene estimates for 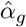 addresses this issue only to a limited extent.

### Variance stabilization transformations implemented in Piccolo

Apart from the *log* transformation as the variance stabilization transformation discussed in the main text and implemented as the default option in `Piccolo`, we also offer the option to apply two other transformations which are described below:

#### Sqrt

For the *sqrt* transformation, 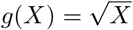. The first order approximations of the mean and variance under this transformation are,

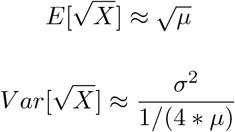

After accounting for sampling depth differences (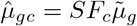and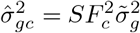) we have,

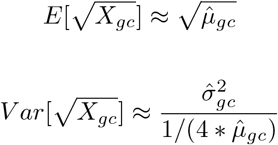

Based on these first order approximations for means and variances under the 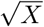 transformation, the residuals are given by,

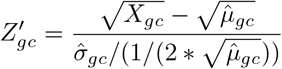

#### Box-Cox Power Law Transform

The Box-Cox transform [57] belongs to the family of power law transformations. Power law transformations are applicable only for positive variables and are indexed by a parameter *λ*, such that for an arbitrary observation *x*, the transformed value is given by *x*^*λ*^. The Box-Cox transformation is a modified power transformation defined as,

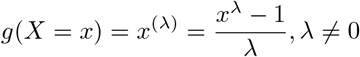

which is a continuous function in *λ* for *x >* 0 (*g*(*X*) = *x*^(*λ*)^ = *log*(*x*), when *λ* = 0). Box-Cox proposed that based on the observations *x*_1_, *x*_2_,…, *x*_*n*_, the appropriate choice of *λ* corresponds to the value that maximizes,

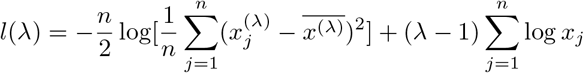

where,

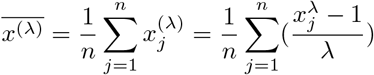

For the Box-Cox transformation, the first-order approximations of the mean and variance are,

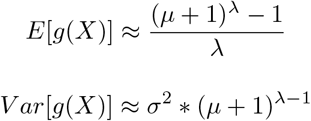

where pseudo-counts of 1 have been added to ensure positive (non-zero) values. After accounting for sampling depth differences (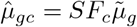 and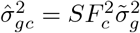) we have,

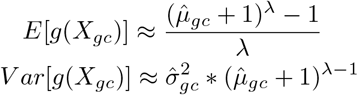

Based on these first-order approximations for means and variances under the Box-Cox transformation, the residuals are given by,

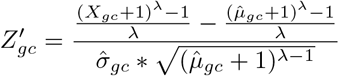

#### LogSF

We also provide the popular `logSF` normalization as an option in `Piccolo`. The normalized counts under this transformation are given by,

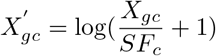

where estimates of *SF*_*c*_ are obtained using stable genes (see eq. (7))

We compared these variance stabilization transformations within Piccolo by applying them to the 100 subsets created using the `Zheng Mix 8eq` data set. The results are shown in Fig. S16. The Box-Cox transform performs the best overall. However, the trade-off is that it is also the most computationally intensive. Nevertheless, we would still recommend users to consider applying it for small or medium sized datasets (10^3^ − 10^5^ cells).

### Differential expression analysis in Piccolo

After applying our normalization to the observed counts, the distribution of the residuals are brought closer to the normal distribution due to the variance stabilization. This makes it possible to employ the two-sample Student’s *t*-test with the null hypothesis that the means of the two samples are the same. Ideally, the test is applicable only if the variances of the two samples can be assumed to be equal. The variance stabilization during normalization ensures that under most circumstances this assumption holds true.

Typically, the preferred test for single-cell differential expression analyses is the Wilcoxon rank-sum test which is a non-parametric alternative to the two-sample *t*-test. A key difference between the two is that while the *t*-test actually tests for location shifts (differences in means) between the samples, the Wilcoxon rank-sum test can be sensitive to shifts in distribution other than a pure location shift.

### Obtaining corrected counts from the *z*-scores

While the clustering and the differential expression analyses are performed using the *z*-scores obtained from `Piccolo`, in some other applications and for the purpose of applying other tools we can also obtain estimates of both the *log*-transformed values as well as the corrected counts. We illustrate how this is done for `Piccolo` normalization (*log*-based variance stabilization).

Recall that,

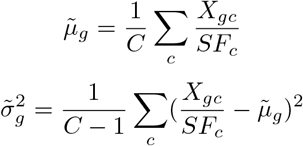

are the respective estimates for the gene mean and variance after accounting for sampling depths. Based on these, the estimates for the *log*-transformed values can be obtained using eq. (16) as,

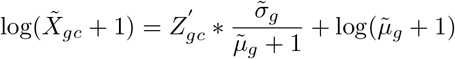

Using these *log*-transformed values, we can then obtain estimates for the corrected counts,

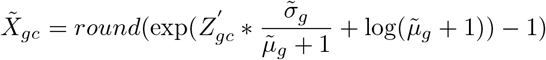

where the rounding ensures that the corrected counts have integer values.

### Batch effect correction

Batch effects are technical confounders that typically stem from a wide array of non-biological sources such as differences in the reagents, the individuals handling and processing the samples, as well as the time of experiment. If not accounted for, batch effect can be misinterpreted as a biological signal and lead to erroneous conclusions. The *z*-score based normalization of `Piccolo` allows for a very simplistic approach to perform batch effect correction based on the fundamental assumption that all the biological conditions are processed in all the batches (Note: It is tempting to apply batch correction even when this assumption is not met, however the conclusions drawn from such analyses will be unreliable).

The basic idea of our batch effect correction method is to identify HVGs independently for each batch, and then seek the HVGs that match between the batches. This ensures that we only retain those genes for downstream analysis that are variable in every batch, and eliminate genes that exhibit variability of counts only within specific batches since the latter likely reflect batch-specific effects. In addition, we identify stable genes across all batches as well as stable genes individually for each batch. We retain only those stable genes in each batch that are also stable across the batches. This ensures that the counts are normalized using size factors estimated with stable genes in each batch that we know are less likely to exhibit batch-specific variation. We then perform the *z*-score normalization independently for each batch. PCA is applied on the composite *z*-scores matrix obtained by combining the *z*-scores matrices from all the batches.

We provide an illustration of the application of our simplistic batch effect correction on a mouse intestinal epithelium data set (`Haber 2017`) that consists of cells obtained in 10 batches [56] (see Fig. S17). Further, we used Splat [38] to simulate a data set using `NIH/3T3` that contained 5 groups of cells, with half the cells in one batch, and the other half simulated to belong to another batch (see Fig. S18). With the help of these examples, we are able to show that the simplistic batch correction approach works well when the assumption that all the conditions are present in every batch is reasonably satisfied.

## Supplementary Tables

**Supplementary Table 1.** Paired Wilcoxon test *p*-values for comparison of the classification metrics between Piccolo and other methods. For each of the 100 datasets created by randomly picking 50% of the cells from the truth-known `Zheng Mix 8eq` data set, we applied Piccolo and the other 3 workflows and then used a kNN based approach to evaluate how well cells belonging to the same cell-types group together. We used paired Wilcoxon test to quantify whether the classification metric values - Macro F1, ARI, AMI - obtained with Piccolo were consistently higher compared to the other approaches (also see Fig. 4B).

**Zheng Mix 8eq - Classification Metrics Comparison**
| Piccolo Compared With | Mean Macro F1 (95% CI) | Paired Wilcoxon Test p-value for Macro F1 | Mean ARI (95% CI) | Paired Wilcoxon Test p-value for ARI | Mean AMI (95% CI) | Paired Wilcoxon Test p-value for AMI |
| --- | --- | --- | --- | --- | --- | --- |
| Piccolo | 0.833 (0.831 - 0.835) | - | 0.780 (0.777 - 0.783) | - | 0.809 (0.808 - 0.811) | - |
| Analytic Pearson | 0.816 (0.815 - 0.818) | 4.33E-18 | 0.768 (0.766 - 0.770) | 1.82E-17 | 0.802 (0.801 - 0.804) | 1.17E-12 |
| Scran LogSF | 0.811 (0.809 - 0.812) | 3.95E-18 | 0.740 (0.738 - 0.743) | 3.95E-18 | 0.761 (0.759 - 0.763) | 3.95E-18 |
| SCTransform v2 | 0.812 (0.810 - 0.814) | 4.60E-18 | 0.760 (0.756 - 0.763) | 4.08E-18 | 0.797 (0.796 - 0.799) | 5.10E-17 |

**Supplementary Table 2.** Paired Wilcoxon test *p*-values for comparison of the classification metrics between Piccolo and other methods where we used the top 3000 HVGs shortlisted by the other methods to compute the residuals using Piccolo normalization. For each of the 100 datasets created by randomly picking 50% of the cells from the truth-known `Zheng Mix 8eq` data set 100 times, we applied Piccolo with the HVGs shortlisted by other methods. We computed the residuals using our normalization method for these HVGs and compared the classification metrics obtained with the corresponding approaches that provided those HVGs (see Fig. 4C). We used paired Wilcoxon test to quantify whether the classification metric values - Macro F1, ARI, AMI - obtained with Piccolo were consistently higher compared to the other approaches.

**Zheng Mix 8eq - Classification Metrics Comparison (Piccolo with other HVGs)**
| Comparison With | Mean Macro F1 (95% CI) | Paired Wilcoxon Test p-value for Macro F1 | Mean ARI (95% CI) | Paired Wilcoxon Test p-value for ARI | Mean AMI (95% CI) | Paired Wilcoxon Test p-value for AMI |
| --- | --- | --- | --- | --- | --- | --- |
| Scran LogSF | 0.811 (0.809 - 0.812) | - | 0.740 (0.738 - 0.743) | - | 0.761 (0.759 - 0.763) | - |
| Piccolo (Scran LogSF) | 0.818 (0.817 - 0.820) | 1.48E-10 | 0.761 (0.759 - 0.763) | 3.95E-18 | 0.784 (0.782 - 0.786) | 3.90E-18 |
| SCTransform v2 | 0.812 (0.810 - 0.814) | - | 0.760 (0.756 - 0.763) | - | 0.797 (0.796 - 0.799) | - |
| Piccolo (SCTransform v2) | 0.827 (0.825 - 0.829) | 6.46E-17 | 0.773 (0.771 - 0.775) | 7.45E-17 | 0.799 (0.797 - 0.800) | 5.74E-02 |

**Supplementary Table 3.** Percentage of genes in each data set that had adjusted *R*^2^ (coefficient of determination) values greater than 0.8. Straight lines were fitted using linear regression through the log-transformed values corresponding to those raw counts that fall within one standard deviation away from the mean expression value. The resultant adjusted *R*^2^ values were recorded and an overall summary in terms of percentage obtained for each data set (see also Fig. S10).

| Data set | Percentage of genes with<br>adjusted $R^2 > 0.8$ |
| --- | --- |
| Svensson 1 | 100% |
| PBMC 33k | 85.1% |
| PBMC r1 (in Drops) | 99.4% |
| NIH/3T3 | 96.5% |
| HEK293T | 97.1% |
| Mouse Cortex r1 (DroNC-seq) | 97% |
| Mouse Lung (Drop-seq) | 97.9% |
| Zheng Mix 8eq | 99.1% |

## Supplementary Figures

**Fig. S1.**
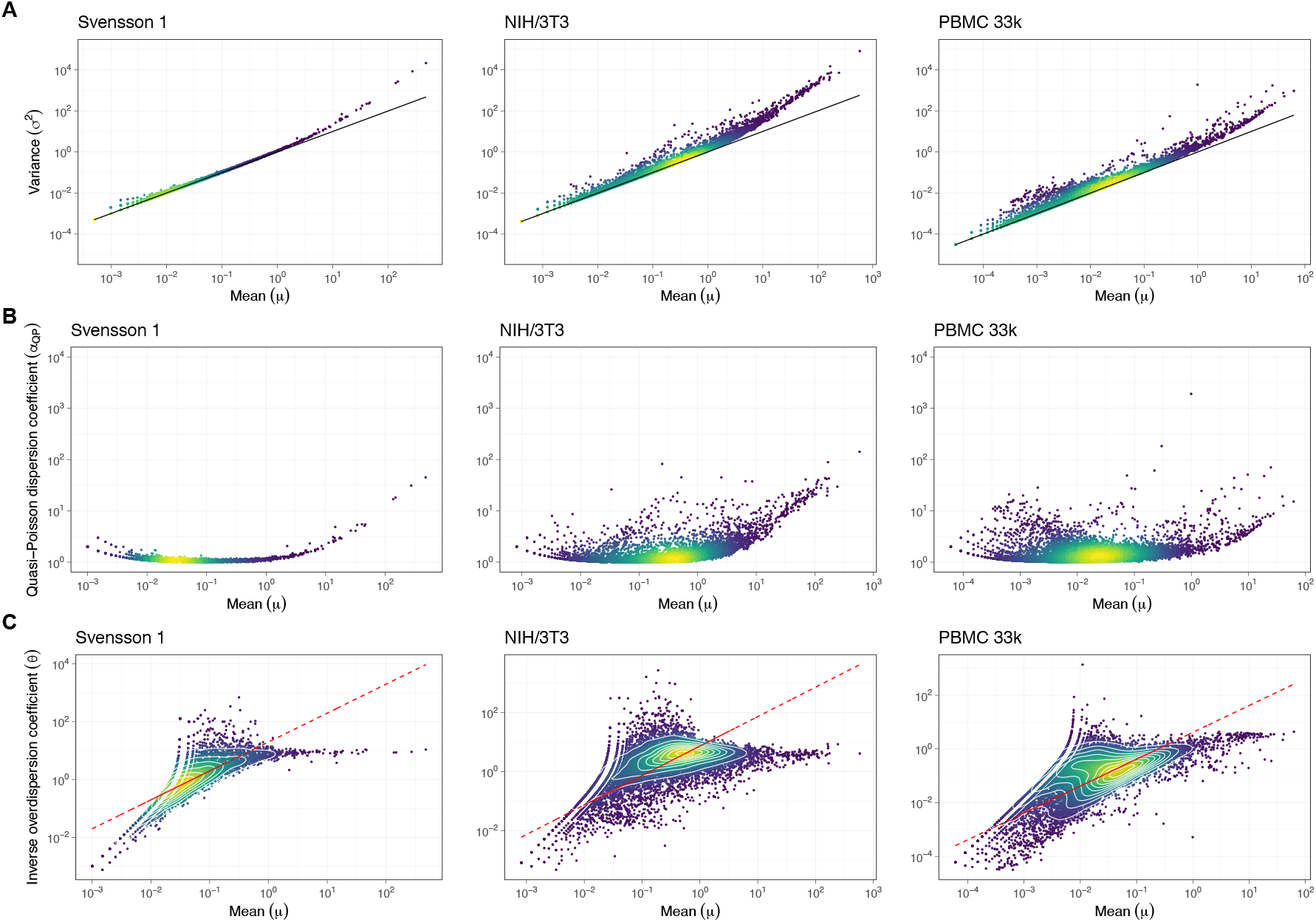
Genes with low mean expression exhibit quasi-Poisson variance. In all the plots, each dot represents a gene and the color of the dots reflect the local point density, with brighter shades (yellow) indicating high density and darker shades (deep blue) indicating low density. **A**. Variance (*σ*^2^) vs mean (*μ*) log-log scatter plots for the Svensson 1 technical control (left panel), NIH/3T3 fibroblast cell line (center panel), and PBMC 33k (right panel) datasets. The solid black line corresponds to the Poisson model (*σ*^2^ = *μ*). For genes with low mean expression levels, the variance can be adequately described by the Poisson model. **B**. Quasi-Poisson dispersion coefficients (*α*_*QP*_) vs mean (*μ*) log-log scatter plots for the Svensson 1 (left panel), NIH/3T3 (center panel), and PBMC 33k (right panel) datasets. *α*_*QP*_ for each gene were estimated from the observed counts using a regression-based approach. **C**. Inverse overdispersion coefficients (*θ*) vs mean (*μ*) log-log scatter plots for the Svensson 1 (left panel), NIH/3T3 (center panel), and PBMC 33k (right panel) datasets. For each gene, *θ* was obtained based on its *α*_*QP*_ using equation. (18). The white curves correspond to the contours lines based on the local point densities. The dashed red line corresponds to estimates of *θ* obtained by fixing *α*_*QP*_ to its value for the gene with the highest point density estimate in the *θ* vs *μ* log-log plot. The monotonically increasing relationship between *θ* and *μ* arises due to the Poisson-like nature of the variance of counts for genes with low expression levels.

**Fig. S2.**
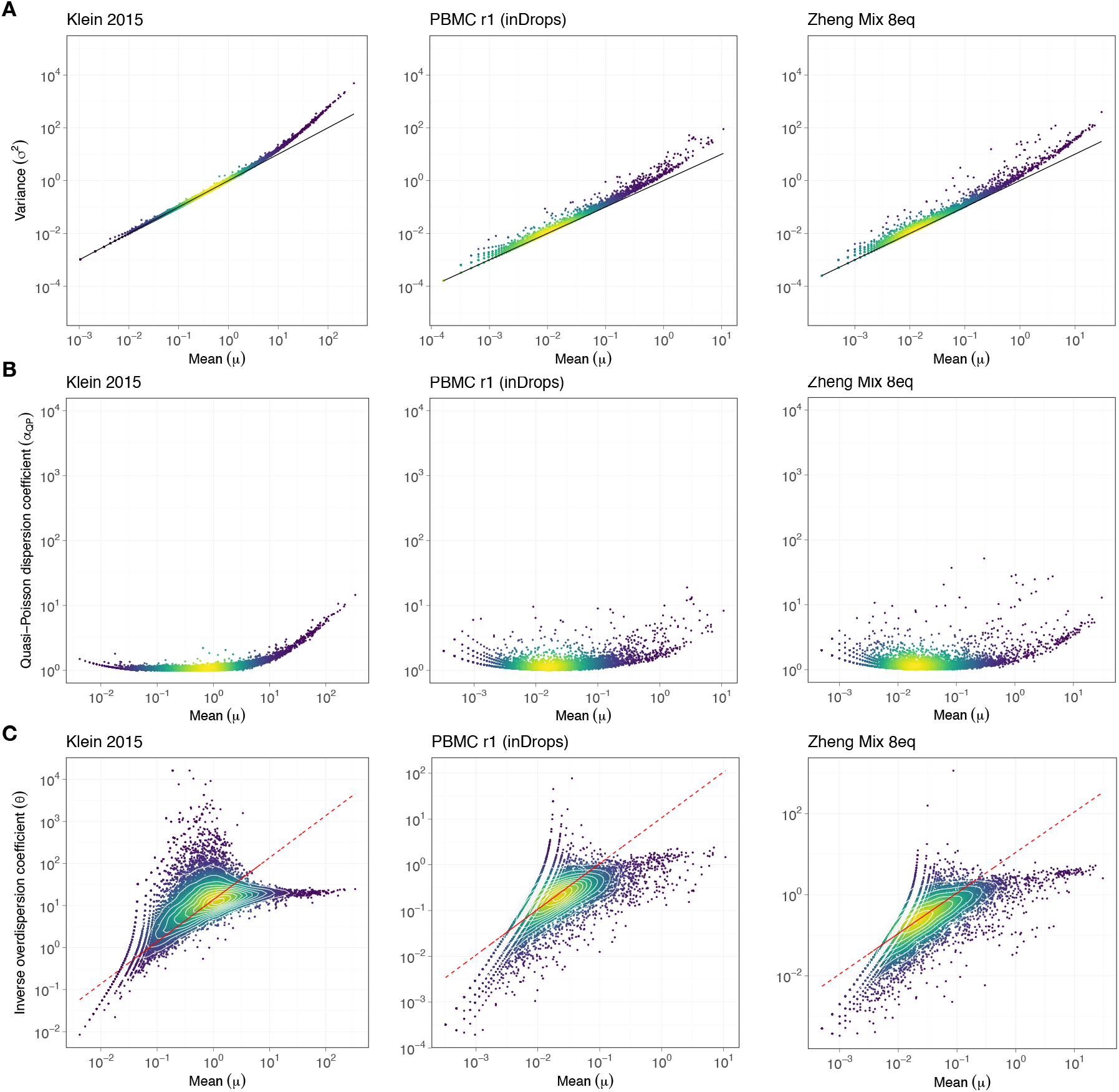
Genes with low mean expression exhibit quasi-Poisson variance. In all the plots, each dot represents a gene and the color of the dots reflect the local point density, with brighter shades (yellow) indicating high density and darker shades (deep blue) indicating low density. **A**. Variance (*σ*^2^) vs mean (*μ*) log-log scatter plots for the Klein 2015 technical control, PBMC r1, and Zheng Mix 8eq datasets (left to right). The solid black line corresponds to the Poisson model (*σ*^2^ = *μ*). For genes with low mean expression levels, the variance is adequately described by the Poisson model. **B**. *α*_*QP*_ vs *μ* log-log scatter plots for the Klein 2015, PBMC r1, and Zheng Mix 8eq datasets. **C**. *θ* vs *μ* log-log scatter plots for the Klein 2015, PBMC r1, and Zheng Mix 8eq datasets. The white curves correspond to the contours lines based on the local point densities. The dashed red line corresponds to estimates of *θ* obtained by fixing *α*_*QP*_ to its value for the gene with the highest point density estimate in the the *θ* vs *μ* log-log plot. The monotonically increasing relationship between *θ* and *μ* for small *μ* indicates quasi-Poisson variance at low mean expression levels.

**Fig. S3.**
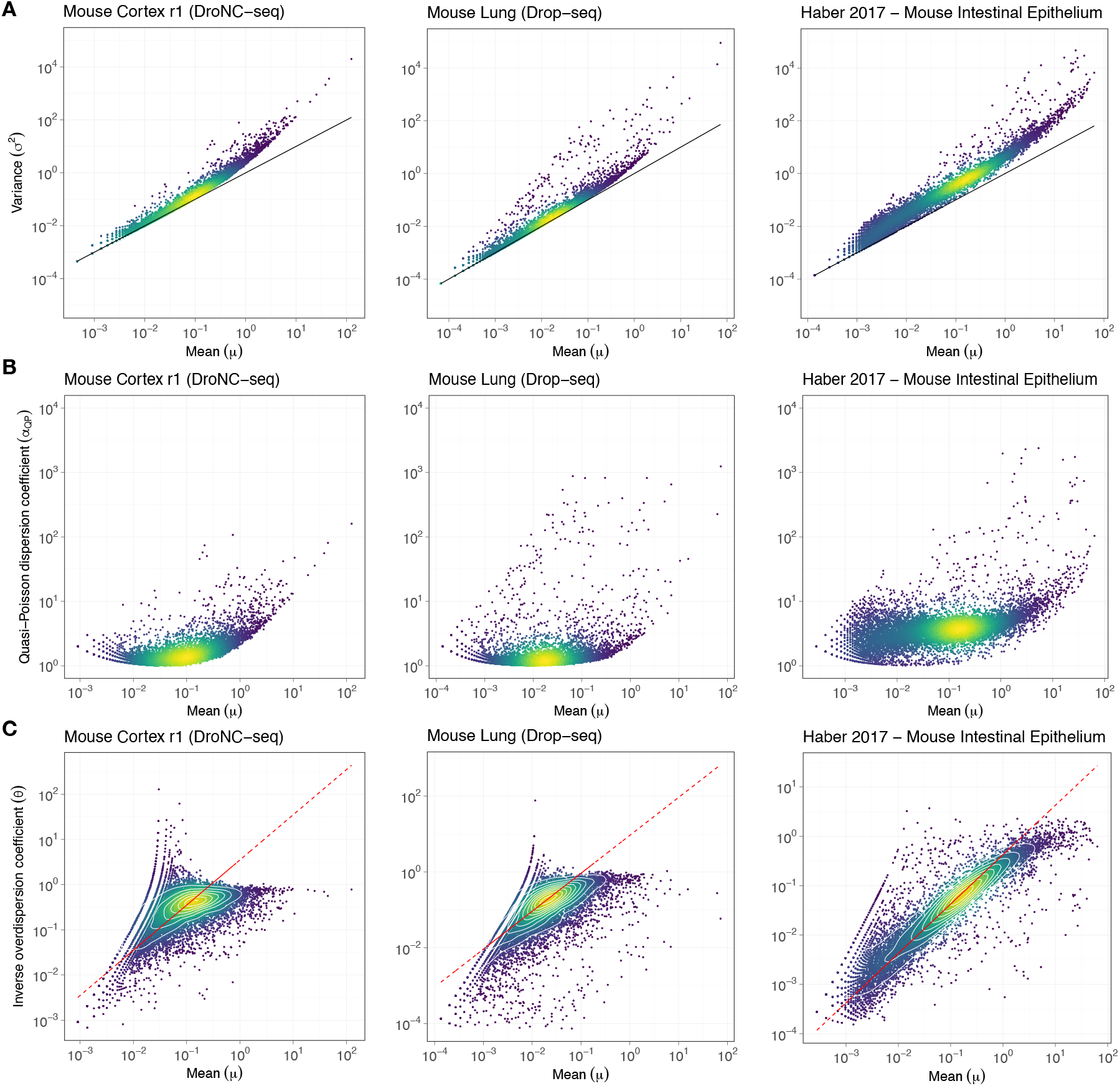
Genes with low mean expression exhibit quasi-Poisson variance. In all the plots, each dot represents a gene and the color of the dots reflect the local point density, with brighter shades (yellow) indicating high density and darker shades (deep blue) indicating low density. **A**. Variance (*σ*^2^) vs mean (*μ*) log-log scatter plots for the Mouse Cortex, Mouse Lung, and Haber 2017 datasets (left to right, in that order). The solid black line corresponds to the Poisson model (*σ*^2^ = *μ*). For genes with low mean expression levels, the variance is adequately described by the Poisson model. **B**. *α*_*QP*_ vs *μ* log-log scatter plots for the Mouse Cortex, Mouse Lung, and Haber 2017 datasets. **C**. *θ* vs *μ* log-log scatter plots for the Mouse Cortex, Mouse Lung, and Haber 2017 datasets. The white curves correspond to the contours lines based on the local point densities. The dashed red line corresponds to estimates of *θ* obtained by fixing *α*_*QP*_ to its value for the gene with the highest point density estimate in the the *θ* vs *μ* log-log plot. The monotonically increasing relationship between *θ* and *μ* for small *μ* indicates quasi-Poisson variance at low mean expression levels.

**Fig. S4.**
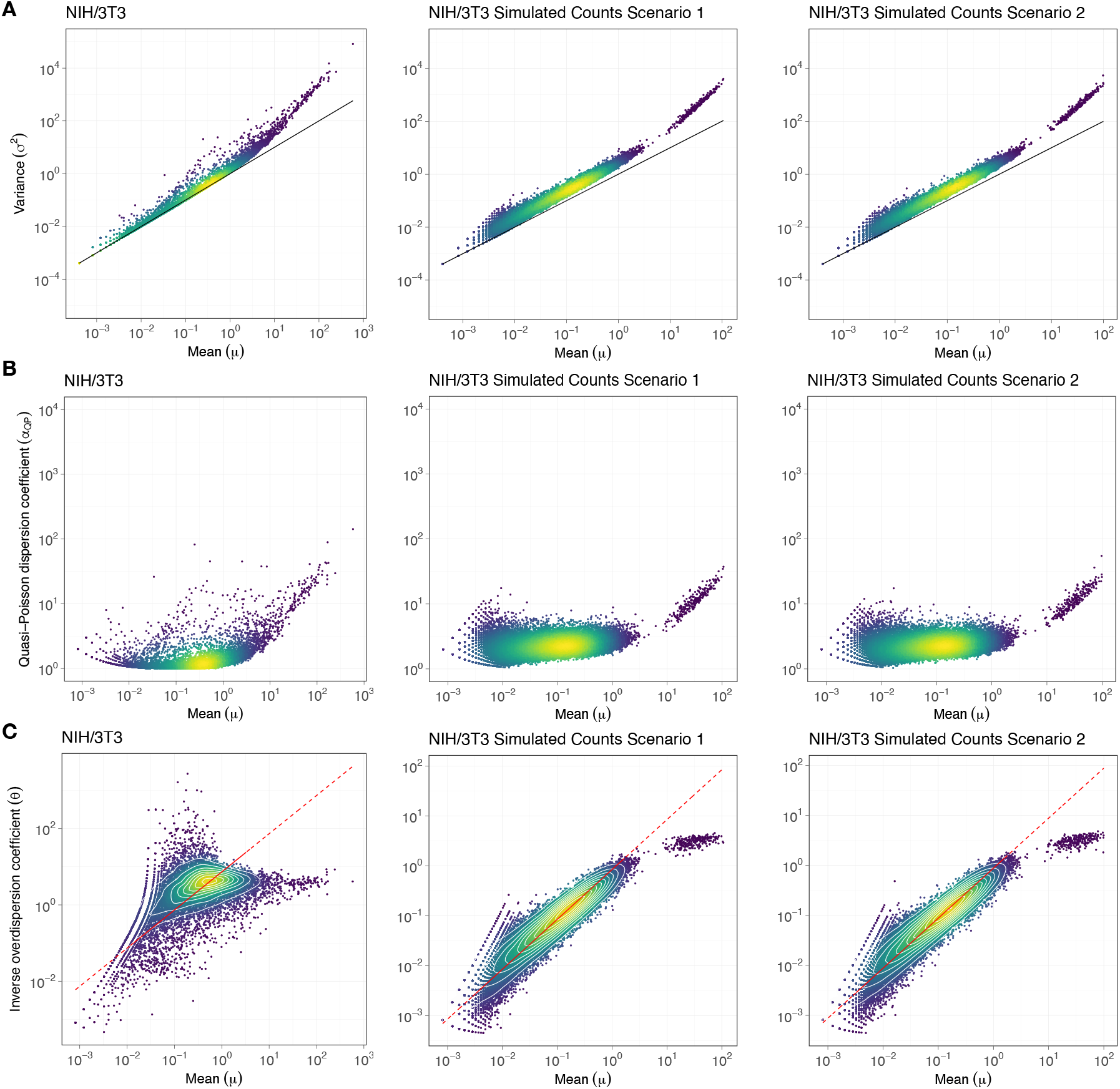
Genes with low mean expression exhibit quasi-Poisson variance. In all the plots, each dot represents a gene and the color of the dots reflect the local point density, with brighter shades (yellow) indicating high density and darker shades (deep blue) indicating low density. **A**. Variance (*σ*^2^) vs mean (*μ*) log-log scatter plots for the NIH/3T3, NIH/3T3 Simulated Counts Scenario 1, and NIH/3T3 Simulated Counts Scenario 2 datasets (left to right). The solid black line corresponds to the Poisson model (*σ*^2^ = *μ*). For genes with low mean expression levels, the variance is adequately described by the Poisson model. **B**. *α*_*QP*_ vs *μ* log-log scatter plots for the NIH/3T3, NIH/3T3 Simulated Counts Scenario 1, and NIH/3T3 Simulated Counts Scenario 2 datasets. **C**. *θ* vs *μ* log-log scatter plots for the NIH/3T3, NIH/3T3 Simulated Counts Scenario 1, and NIH/3T3 Simulated Counts Scenario 2 datasets. The white curves correspond to the contours lines based on the local point densities. The dashed red line corresponds to estimates of *θ* obtained by fixing *α*_*QP*_ to its value for the gene with the highest point density estimate in the the *θ* vs *μ* log-log plot. The Splat [38] simulated counts datasets exhibit pronounced quasi-Poisson variance (monotonically increasing relationship between *θ* and *μ* for small *μ*) compared to the original data set.

**Fig. S5.**
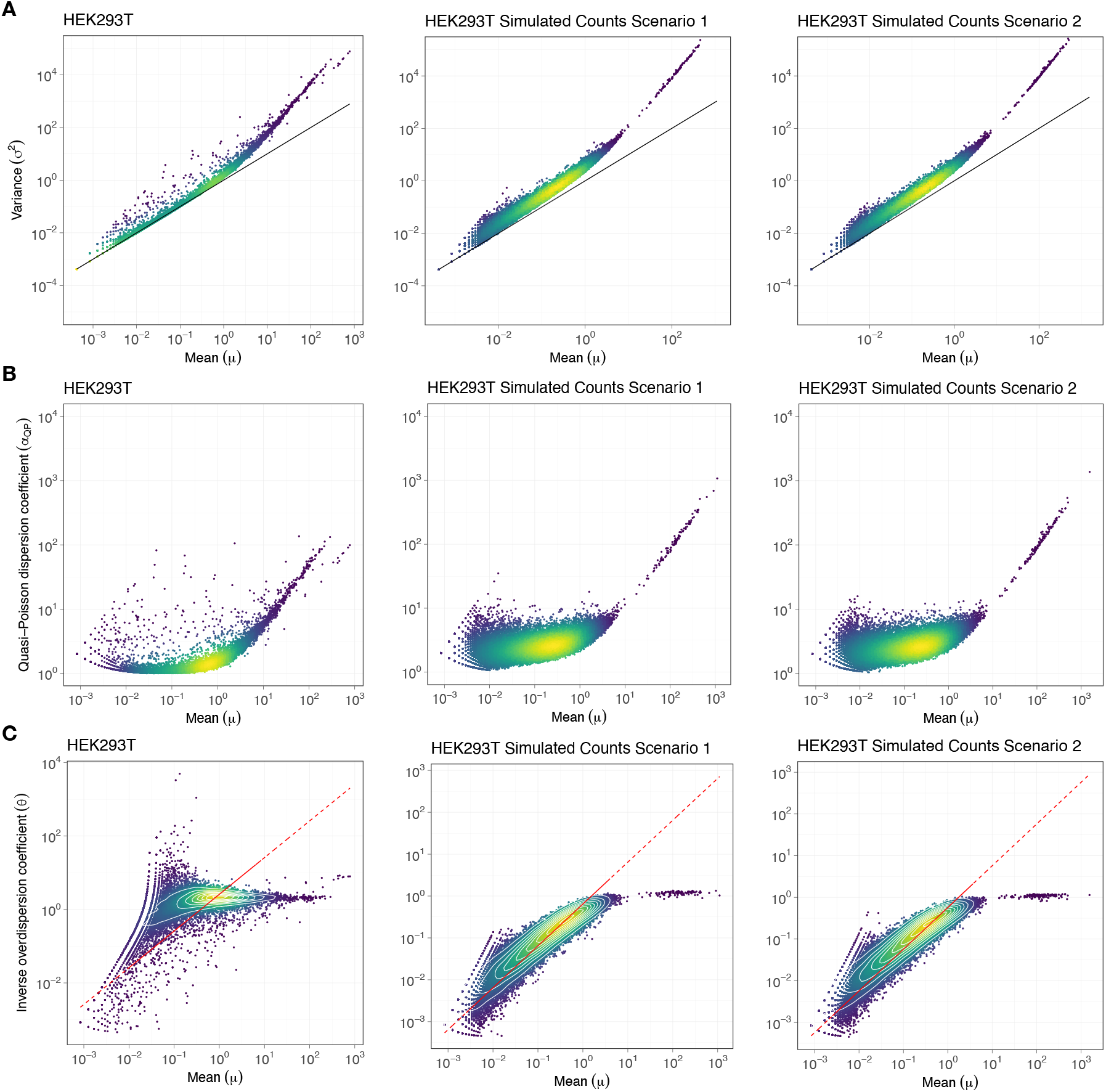
Genes with low mean expression exhibit quasi-Poisson variance. In all the plots, each dot represents a gene and the color of the dots reflect the local point density, with brighter shades (yellow) indicating high density and darker shades (deep blue) indicating low density. **A**. Variance (*σ*^2^) vs mean (*μ*) log-log scatter plots for the HEK293T, HEK293T Simulated Counts Scenario 1, and HEK293T Simulated Counts Scenario 2 datasets (left to right). The solid black line corresponds to the Poisson model (*σ*^2^ = *μ*). For genes with low mean expression levels, the variance is adequately described by the Poisson model. **B**. *α*_*QP*_ vs *μ* log-log scatter plots for the HEK293T, HEK293T Simulated Counts Scenario 1, and HEK293T Simulated Counts Scenario 2 datasets. **C**. *θ* vs *μ* log-log scatter plots for the HEK293T, HEK293T Simulated Counts Scenario 1, and HEK293T Simulated Counts Scenario 2 datasets. The white curves correspond to the contours lines based on the local point densities. The dashed red line corresponds to estimates of *θ* obtained by fixing *α*_*QP*_ to its value for the gene with the highest point density estimate in the the *θ* vs *μ* log-log plot. The Splat [38] simulated counts datasets exhibit pronounced quasi-Poisson variance (monotonically increasing relationship between *θ* and *μ* for small *μ*) compared to the original data set.

**Fig. S6.**
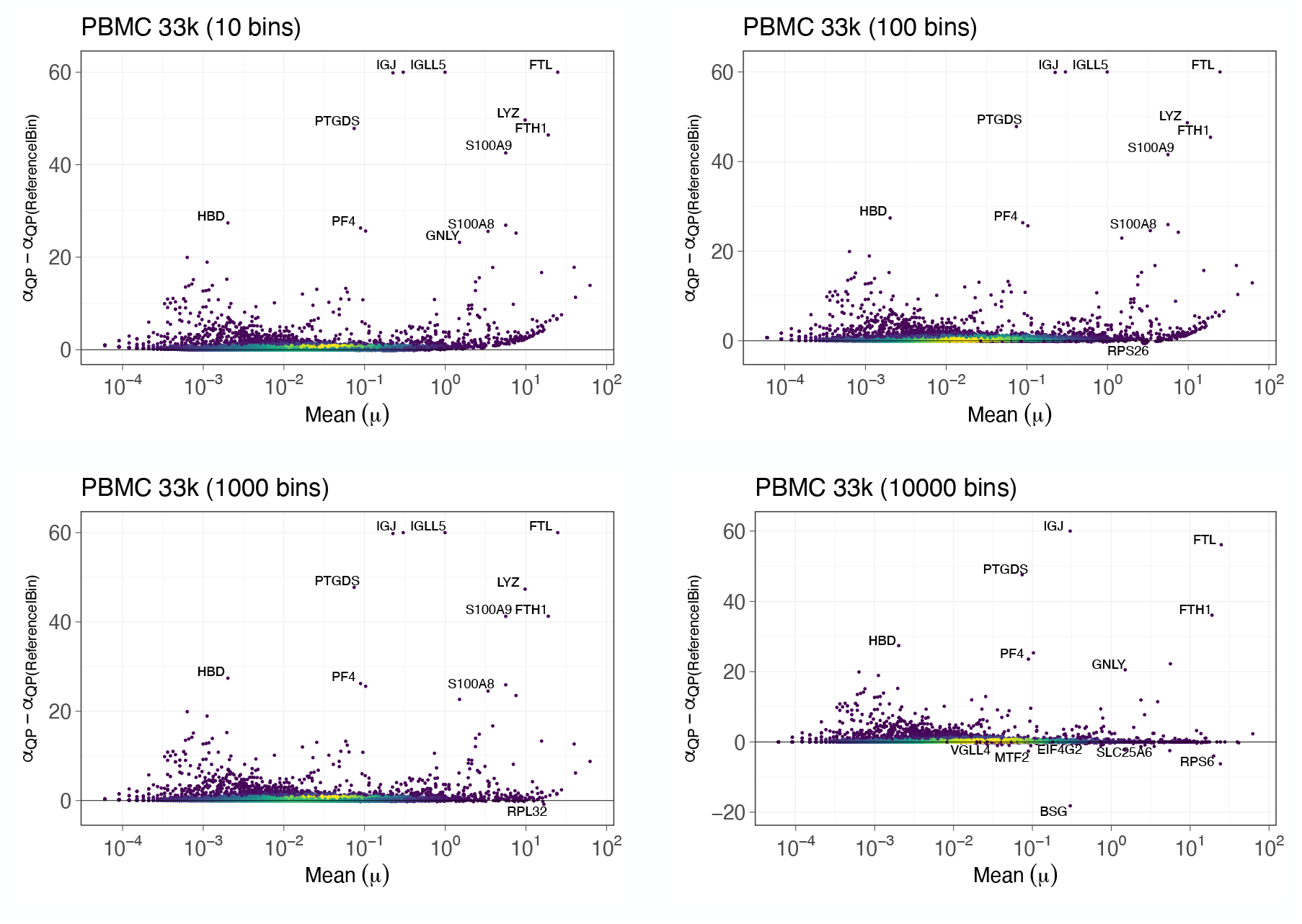
The number of bins affect which genes are shortlisted as HVGs, particularly among genes with high mean expression levels. In all the panels, each dot is a gene and is colored based on the local density of the dots, with dots in higher density regions shown in brighter shades (yellow) and lower density regions shown in darker shades (dark blue). Each panel depicts *α*_*QP*_−*α*_*QP* (*Reference*|*Bin*)_ vs mean (*μ*) linear-log scatter plots for genes that exhibited over-dispersion with respect to the Poisson (*σ*^2^ *> μ*) for `PBMC 33k` corresponding to different choices for number of bins in the feature selection step. Fewer bins imply that there are more genes per bin and this leads to underestimation of the reference quantile especially for the bin that contains genes with the largest mean expression levels since their *α*_*QP*_ vary the most with *μ*. Note the pronounced increase of *α*_*QP*_ −*α*_*QP* (*Reference*|*Bin*)_ with *μ* in the top left panel for high mean expression genes. This will lead to a bias towards selection of genes with high mean expression as HVGs. Increasing the number of bins leads to a reduction in this bias (see bottom left panel). However, if we keep on increasing the number of bins there will be very few genes per bin. Since we effectively assume that in every bin there is a stable gene we would end up concluding for a significant proportion of genes that these genes are stable even if they actually exhibit biological variability. In practice, setting the number of bins to its default value of 1000 leads to approximately 10-15 genes per bin with the gene with the smallest or the second smallest value within the bin providing the *α*_*QP* (*Reference*|*Bin*)_ value for that bin.

**Fig. S7.**
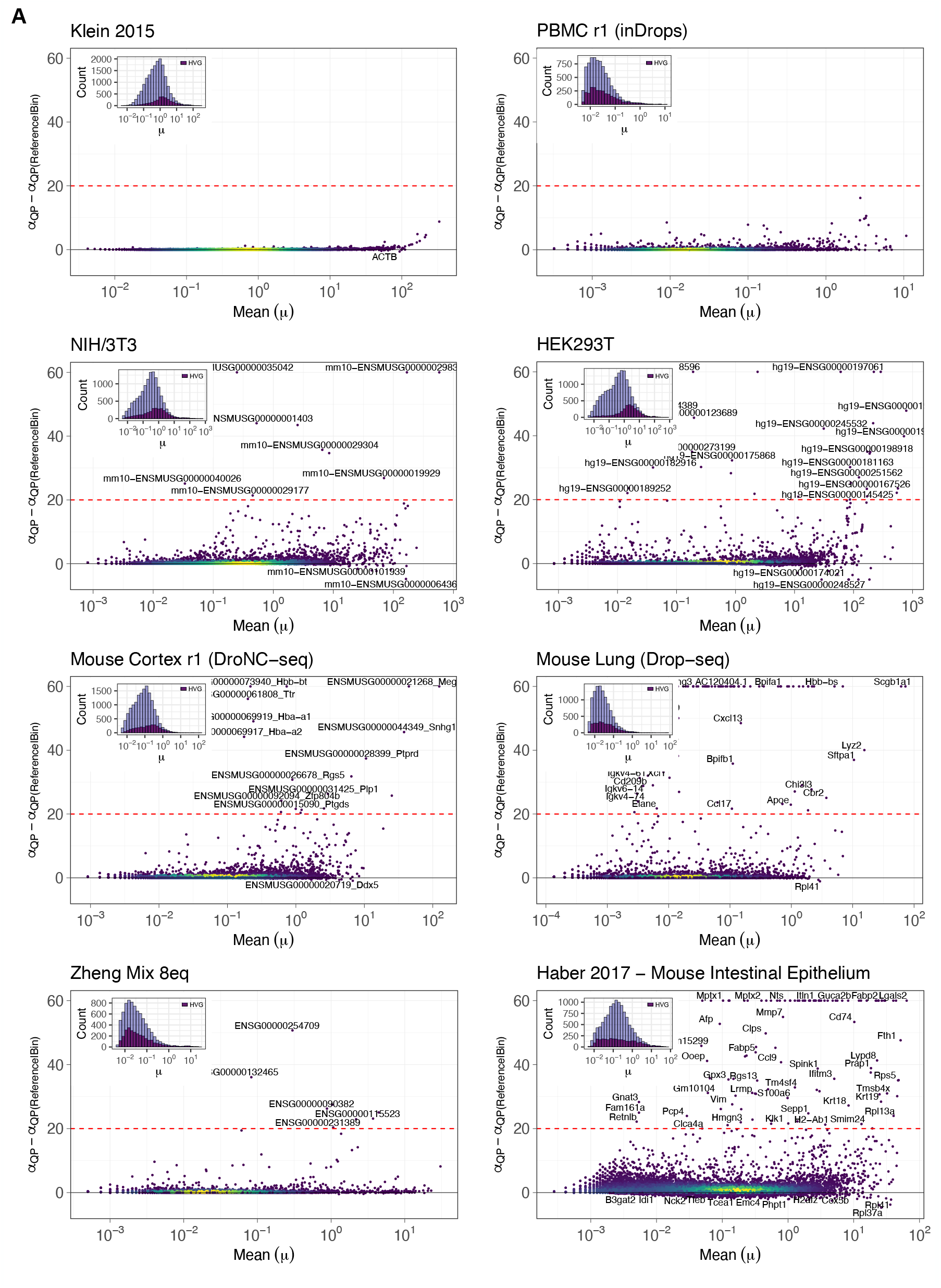
Feature selection using bin-based approach does not lead to a preferential selection of genes based on their mean expression level. In all the panels, each dot is a gene and is colored based on the local density of the dots, with dots in higher density regions shown in brighter shades (yellow) and lower density regions shown in darker shades (dark blue). **A**. *α*_*QP*_−*α*_*QP* (*Reference*|*Bin*)_ vs mean (*μ*) linear-log scatter plot for genes that exhibited over-dispersion with respect to the Poisson (*σ*^2^ *> μ*) for 8 datasets (the panels are titled with the names of the respective datasets). The dashed red horizontal line illustrates the threshold to shortlist HVGs. Genes with *α*_*QP*_ − *α*_*QP* (*Reference*|*Bin*)_ *>* 20 in this case are shortlisted as HVGs. On the other hand, genes with *α*_*QP*_ − *α*_*QP* (*Reference*|*Bin*)_ *<* 0 are shortlisted as stable genes. Inset in top left corner - light colored histogram corresponding to all genes with *α*_*QP*_ *>* 1, and dark colored histogram corresponding to the top 3000 HVGs.

**Fig. S8.**
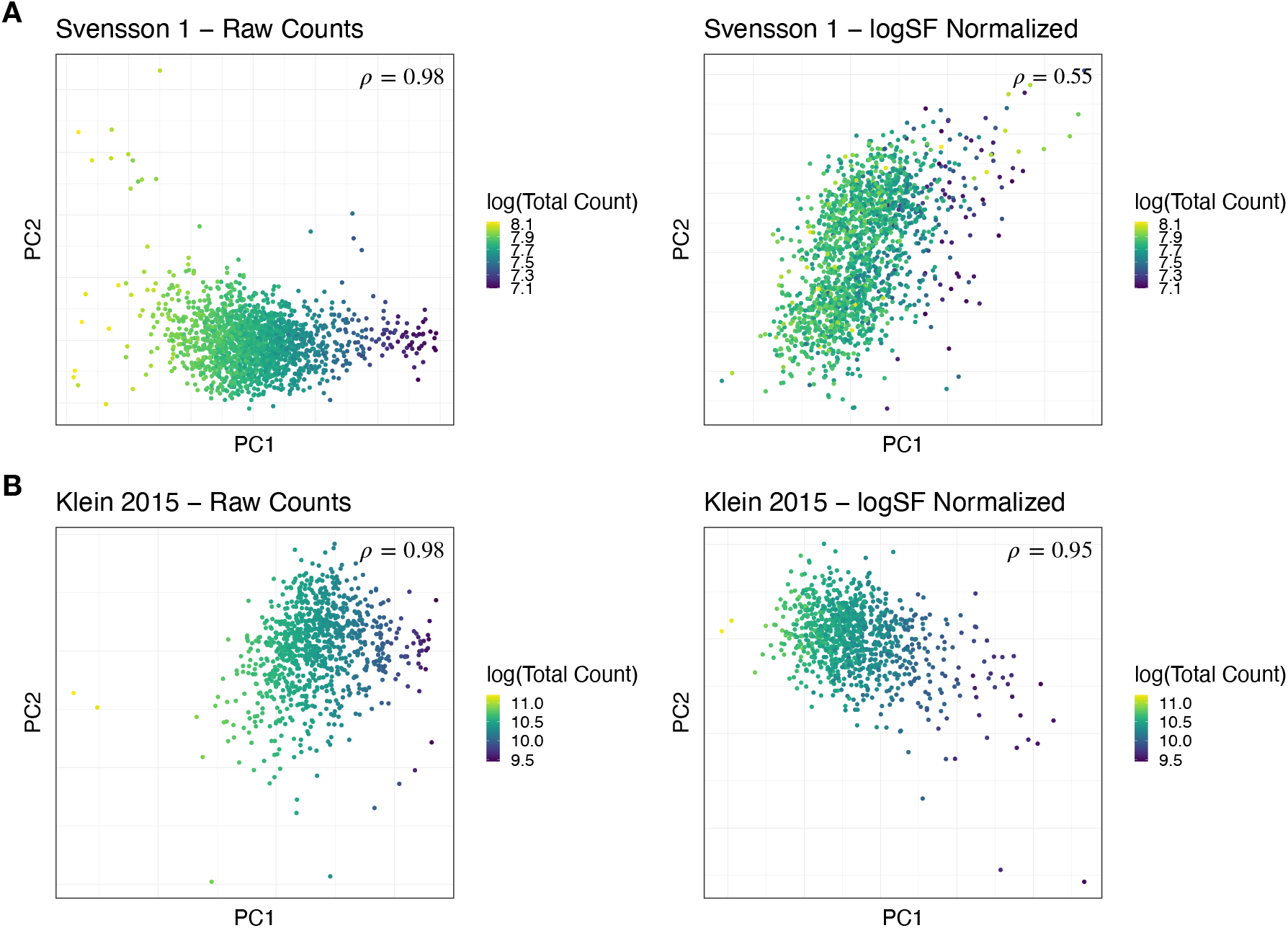
LogSF normalization does not significantly reduce the impact of sampling depth (total count) differences. **A**. 2D scatter plots based on the first 2 PCs of the Svensson 1 technical control data set. Each dot is a cell and is colored according to the total counts; brighter shades (yellow) correspond to larger total counts and darker shades (deep blue) correspond to smaller total counts. The left panel shows the plot for the raw counts, while the right panel shows the plot for the logSF transformed counts. The coefficient (*ρ*) in the top-right corner of the panels shows the canonical correlation coefficient between the total counts per cell and the top 5 PCs. Smaller values of *ρ* indicate lesser correlation between the total counts and the top 5 PCs, thus indicating that sampling depth differences have been reduced more effectively by the normalization. LogSF normalization does not reduce the impact of sampling depth on the overall variation as evaluated through PCA since the *ρ* has not been reduced significantly. **B**. 2D scatter plots based on the first 2 PCs of the Klein 2015 technical control data set. The left panel shows the plot for the raw counts, while the right panel shows the plot for the logSF transformed counts. The coefficient (*ρ*) in the top-right corner of the panels shows the canonical correlation coefficient between the total counts per cell and the top 5 PCs, and clearly indicates that even after logSF normalization the sampling depth differences are still contributing significantly to the overall variance in the data set.

**Fig. S9.**
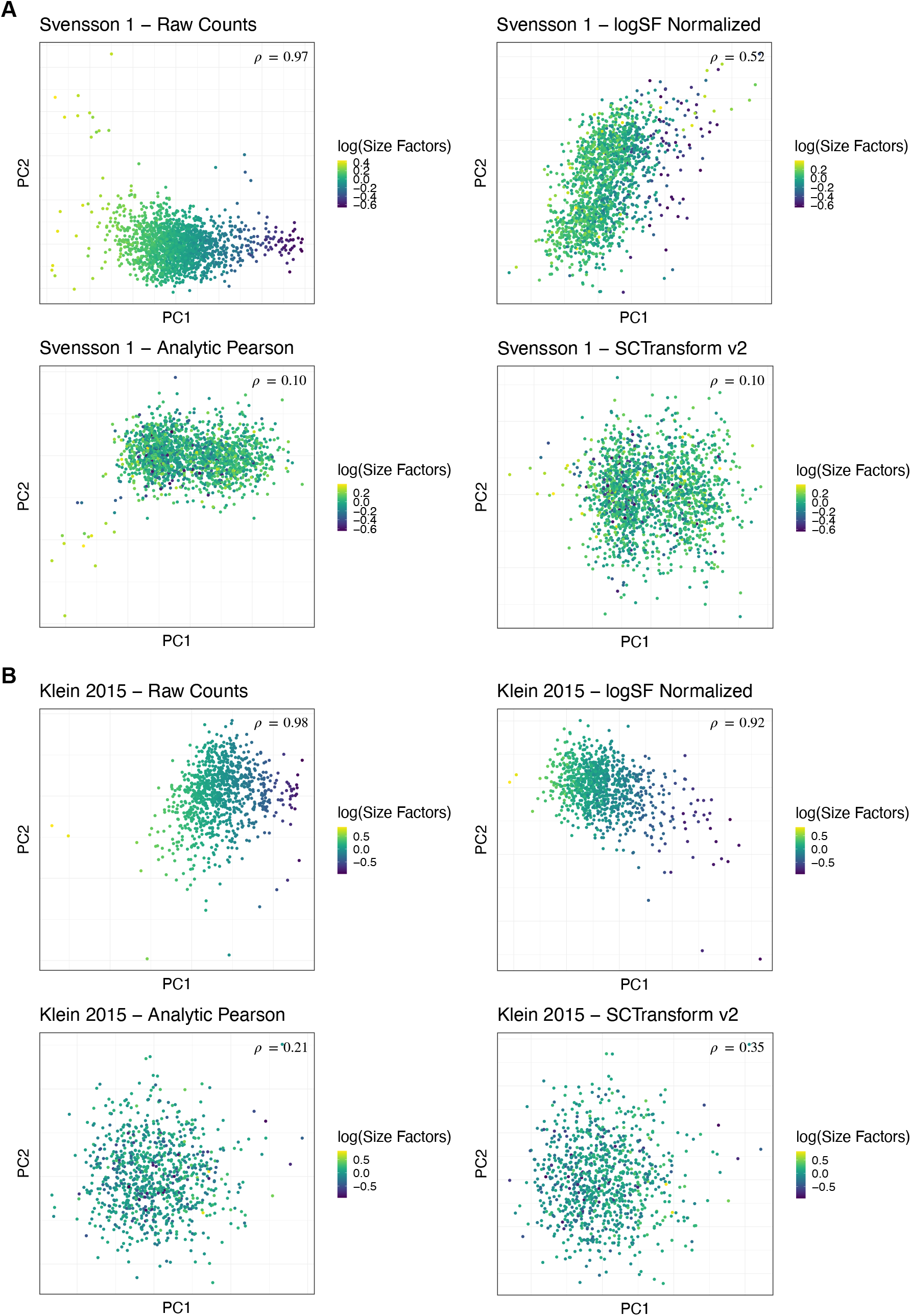
Residuals-based approaches reduce the impact of sampling depth differences more effectively than logSF normalization. **A**. 2D scatter plots based on the first 2 PCs of the Svensson 1 technical control data set. Each dot is a cell and is colored according to the size factors; brighter shades (yellow) correspond to larger size factors and darker shades (deep blue) correspond to smaller size factors. The top-left panel shows the plot for the raw counts, while the top-right panel shows the plot for the logSF transformed counts, the bottom 2 panels show the plots for the residuals obtained using Analytic Pearson method and SCTransform v2 respectively. The coefficient (*ρ*) in the top-right corner of the panels shows the canonical correlation coefficient between the size factors and the top 5 PCs. Smaller values of *ρ* indicate that the impact of the sampling depth differences has been reduced more effectively by the normalization. The values of *ρ* for the residuals-based methods are significantly smaller than the value obtained using logSF normalization. **B**. Same as in **A**. except this is for the Klein 2015 technical control data set. Once again, the residuals-based methods lead to much lesser correlation between the top 5 principal components and the size factors.

**Fig. S10.**
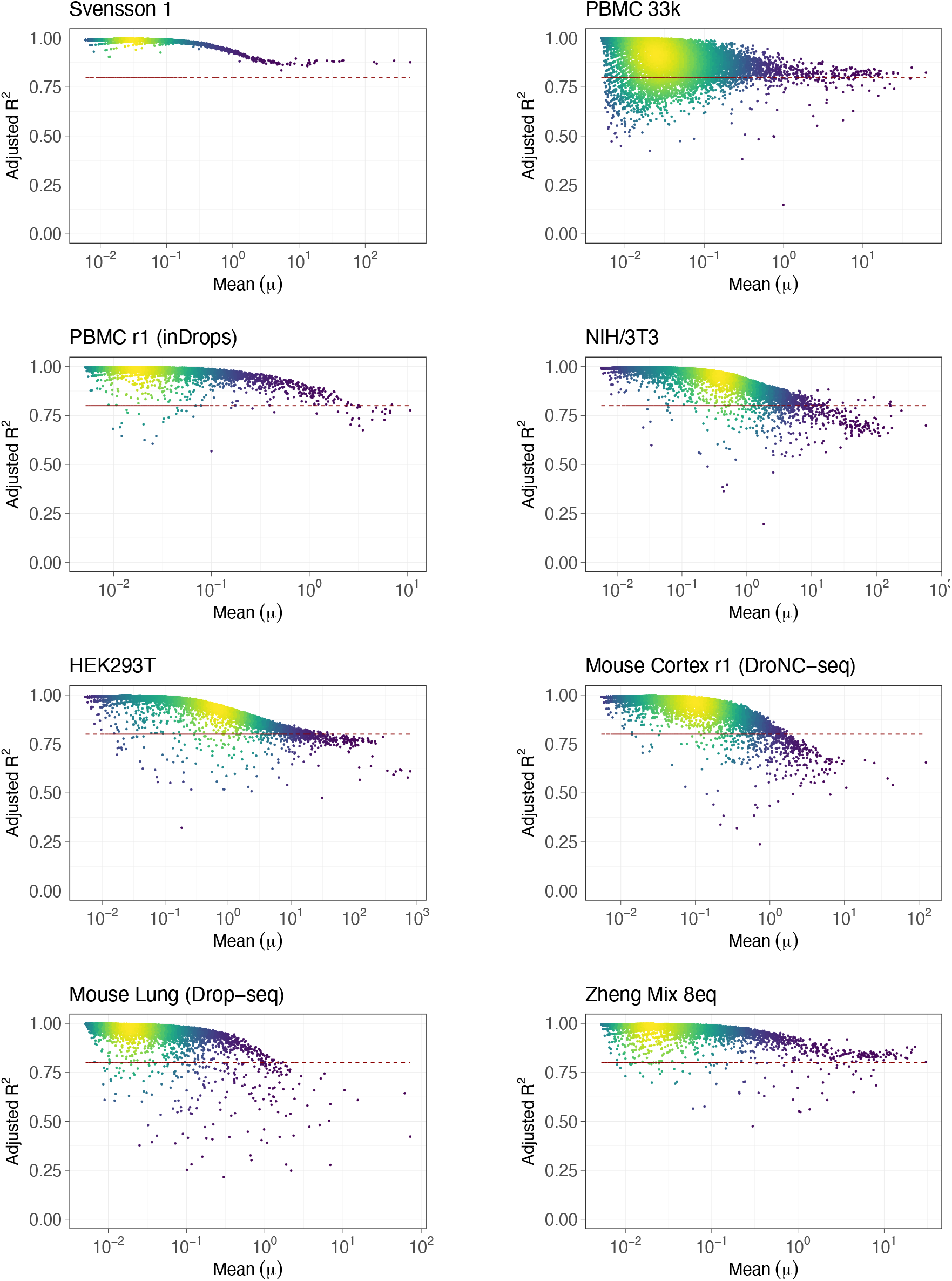
First-order approximation for the transformed counts under variance stabilization transformation applies quite well for majority of the genes. For each gene, we fitted a straight line using linear regression through the log transformed values corresponding to those raw counts that fall within one standard deviation away from the mean expression value. Here we show the resultant adjusted *R*^2^ (coefficient of determination) values for the fitted straight lines for each gene. For most of the genes (especially those with low expression) the adjusted *R*^2^ are close to 1 indicating that the first-order approximation is valid for the transformed values of these genes.

**Fig. S11.**
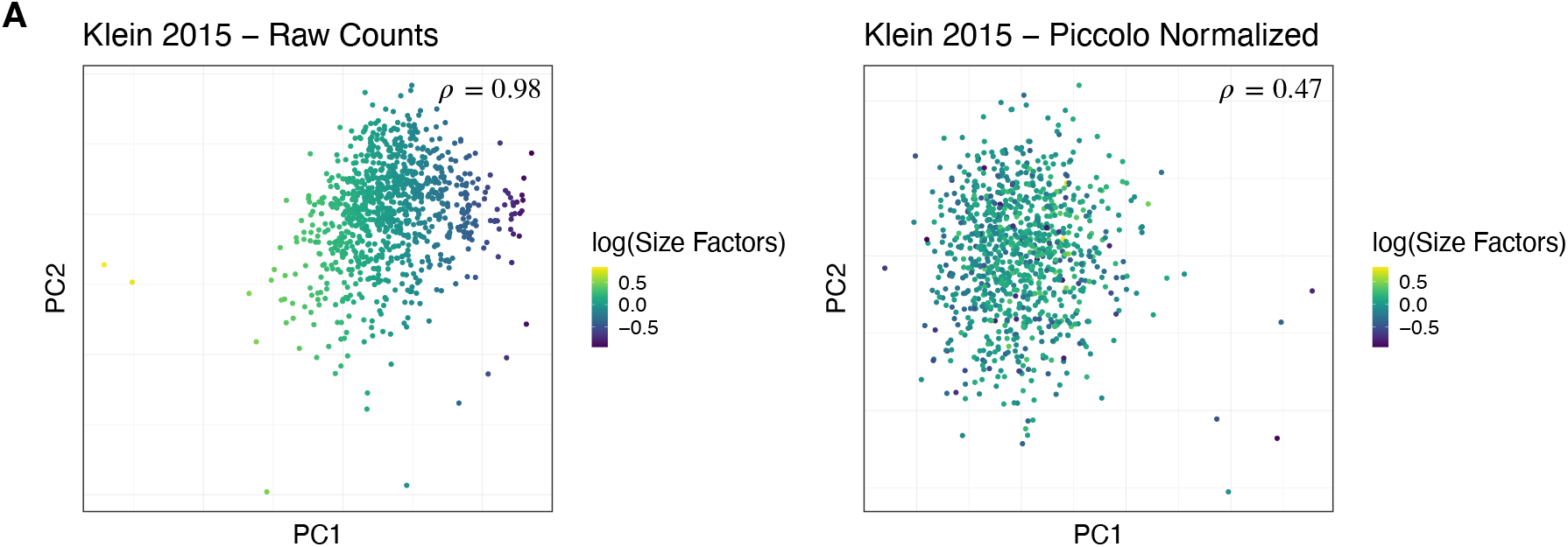
Piccolo normalization reduces the impact of sampling depth differences. **A**. 2D scatter plots based on the first 2 PCs of the Klein 2015 technical control data set. Each dot is a cell and is colored according to the size factors; brighter shades (yellow) correspond to larger size factors and darker shades (deep blue) correspond to smaller size factors. The left panel shows the plot for the raw counts, while the right panel shows the plot for the residuals obtained using Piccolo normalization. The coefficient (*ρ*) in the top-right corner of the panels shows the canonical correlation coefficient between the size factors and the top 5 PCs. The value of *ρ* for the residuals obtained with Piccolo is significantly smaller than the value for the raw counts.

**Fig. S12.**
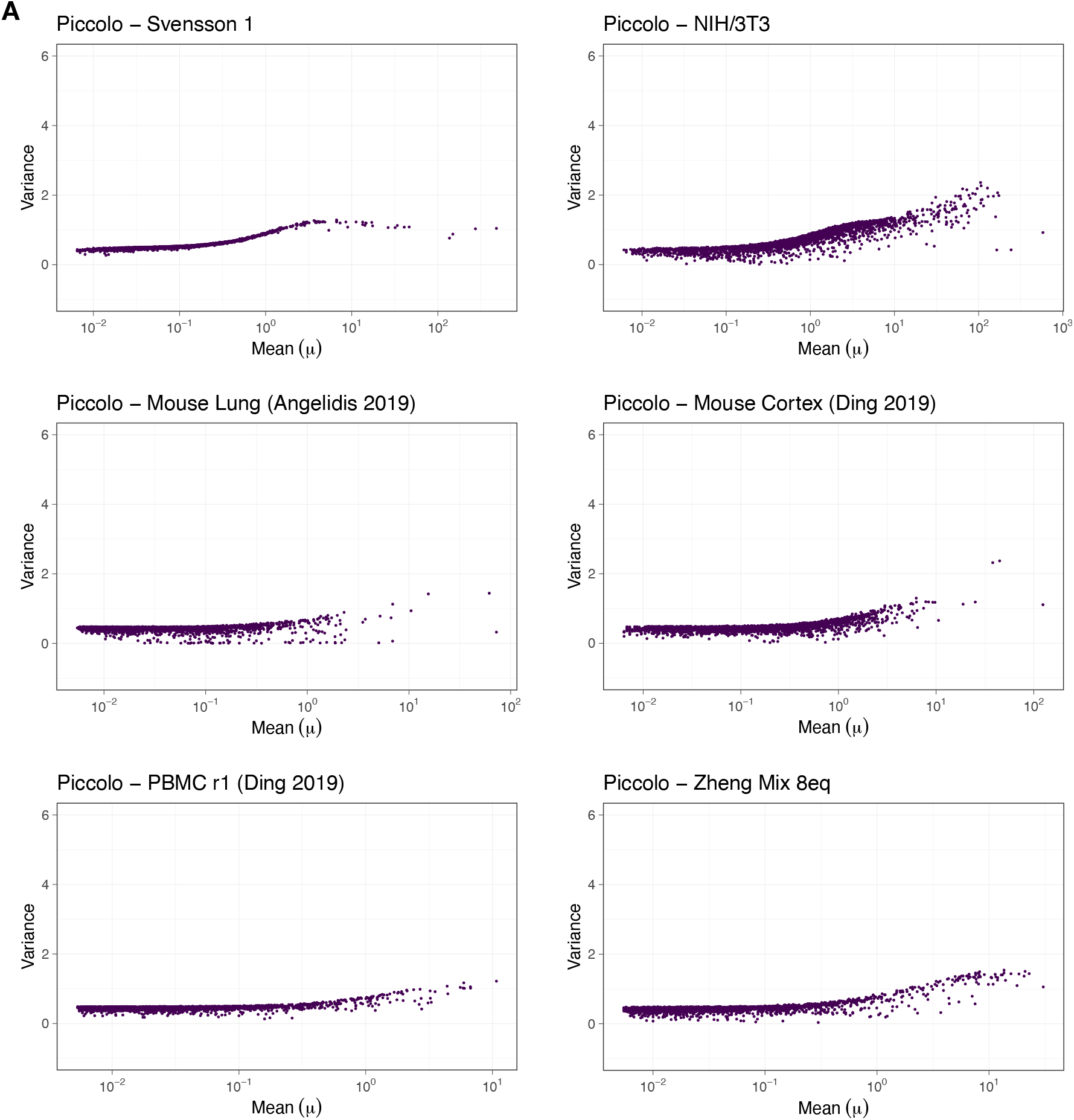
Piccolo normalization ensures variance stabilization. **A**. Residual variance vs mean (*μ*) linear-log scatter plots for the top 3000 HVGs of 6 datasets after applying Piccolo normalization. The residuals obtained using Piccolo do not exhibit significant deviations away from 1 unlike residuals derived from Analytic Pearson and SCTransform.

**Fig. S13.**
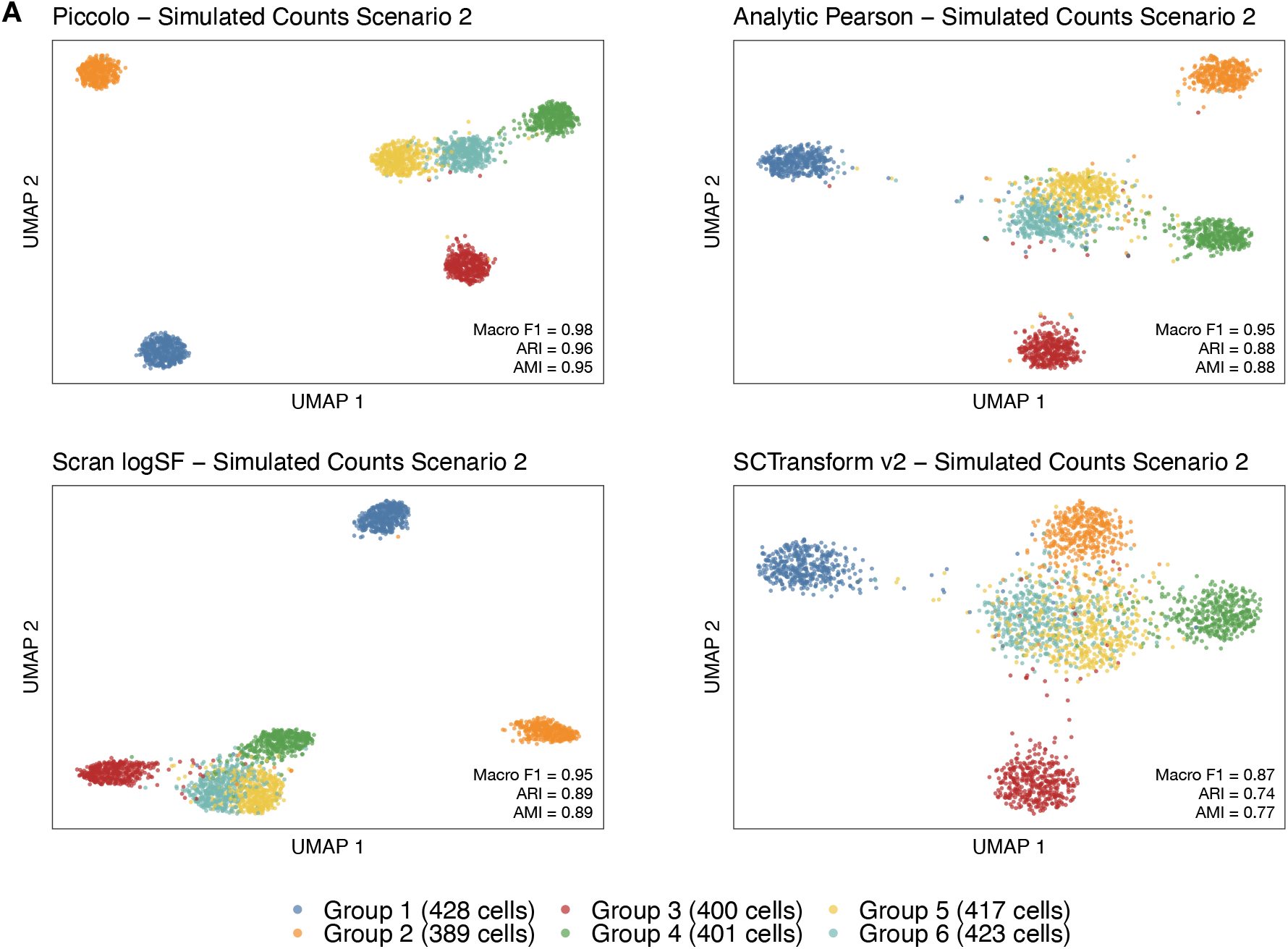
Piccolo enables robust clustering of cell groups, including cell groups with few differentially expressed genes. **A**. 2D UMAP plots after using respective normalization methods for the top 3000 HVGs for the Simulated Counts Scenario 2 data set - 6 groups with approximately same number of cells per group but with different number of DE genes per group. The simulated counts were generated by Splat [38] using the NIH/3T3 data set. Dots represent cells and are colored using the known cell-type labels (legend at the bottom of the panel; actual numbers of cells belonging to each group are specified in parentheses). Clustering metrics based on comparisons between predicted cell labels and known cell labels are listed in the bottom-right corner in each panel. With Piccolo, cells belonging to the 6 groups cluster distinctly and can be distinguished better compared to other normalization methods.

**Fig. S14.**
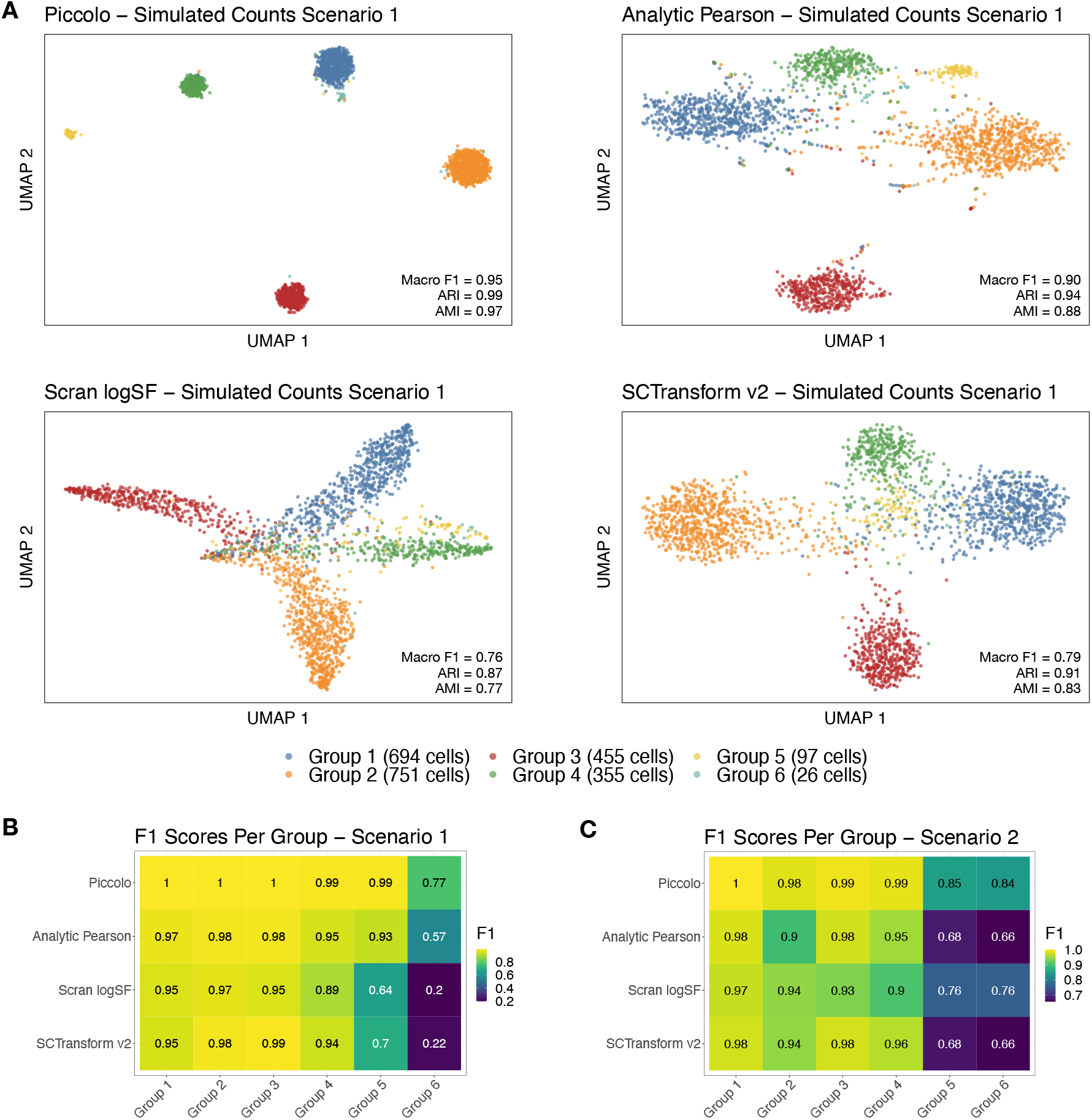
Piccolo enables identification of clusters even for groups containing small numbers of cells. **A**. 2D UMAP plots after using respective normalization methods for the top 3000 HVGs for the Simulated Counts Scenario 1 data set. The simulated counts were generated by Splat [38] using the HEK293T data set. Dots represent cells and are colored using the known cell-type labels (legend at the bottom of the panel; actual numbers of cells belonging to each group are specified in parentheses). Clustering metrics based on comparisons between predicted cell labels and known cell labels are listed in the bottom-right corner in each panel. Cells belonging to the 6 groups can be distinguished using Piccolo. **B**. Heatmap showing the F1 scores per group for each of the normalization methods applied to Simulated Counts Scenario 1. The tiles of the heatmap are colored according to the F1 values with larger F1 scores corresponding to brighter shades (yellow), and lower F1 scores corresponding to darker shades (deep blue). Piccolo has the highest F1 scores for all 6 groups. **C**. Heatmap of F1 scores per group for the Simulated Counts Scenario 2 data set (see Fig. S12). Piccolo returns the highest F1 scores for all 6 groups.

**Fig. S15.**
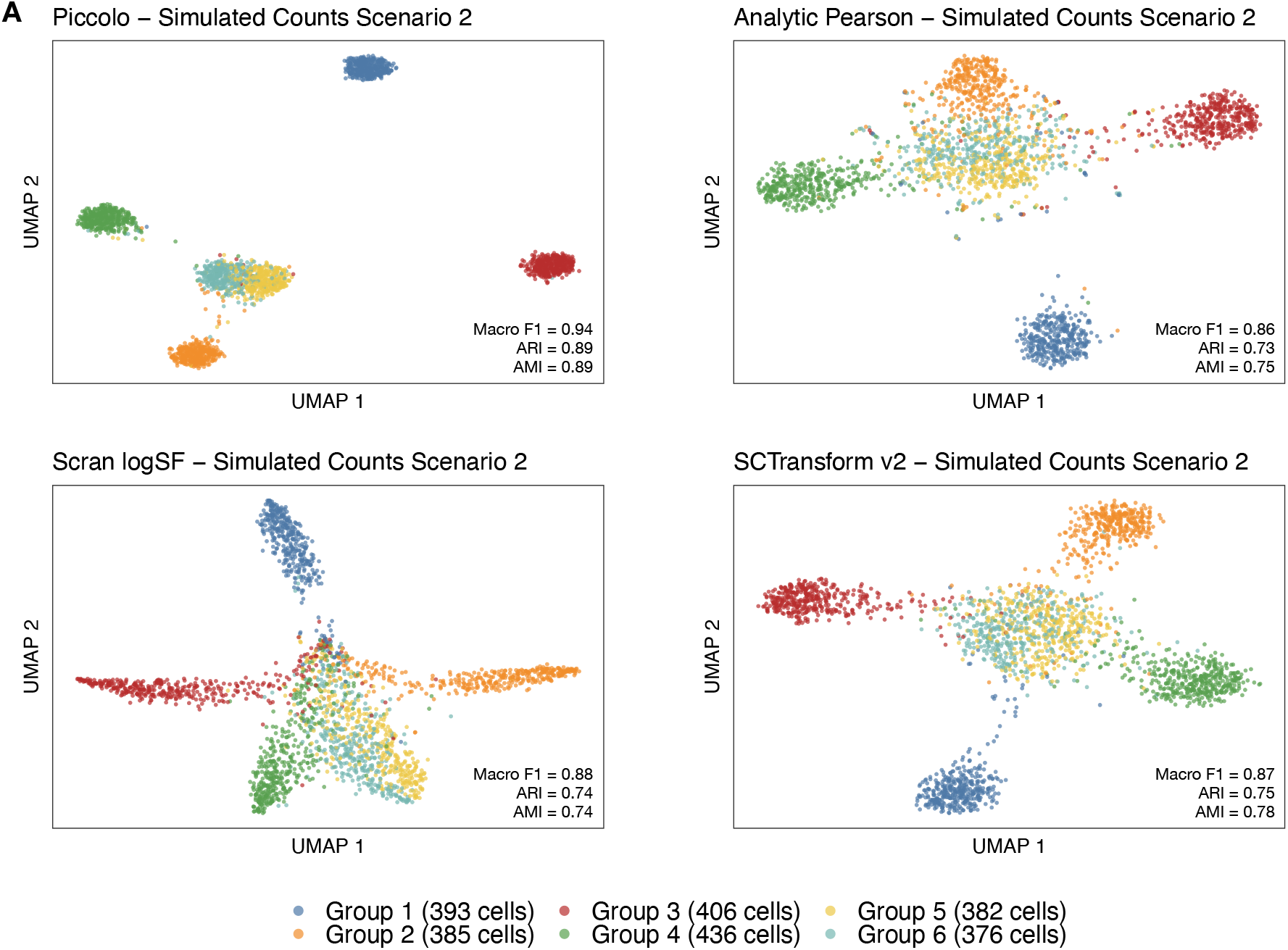
Piccolo enables robust clustering of cell groups, including cell groups with few differentially expressed genes. **A**. 2D UMAP plots after using respective normalization methods for the top 3000 HVGs for the simulated counts data set - Simulated Counts Scenario 2 (6 groups with approximately same number of cells per group but with different number of DE genes per group). The simulated counts were generated by Splat [38] using the HEK293T data set. Dots represent cells and are colored using the known cell-type labels (legend at the bottom of the panel; actual numbers of cells belonging to each group are specified in parentheses). Clustering metrics based on comparisons between predicted cell labels and known cell labels are listed in the bottom-right corner in each panel. With Piccolo, cells belonging to the 6 groups cluster distinctly and can be distinguished better compared to other normalization methods.

**Fig. S16.**
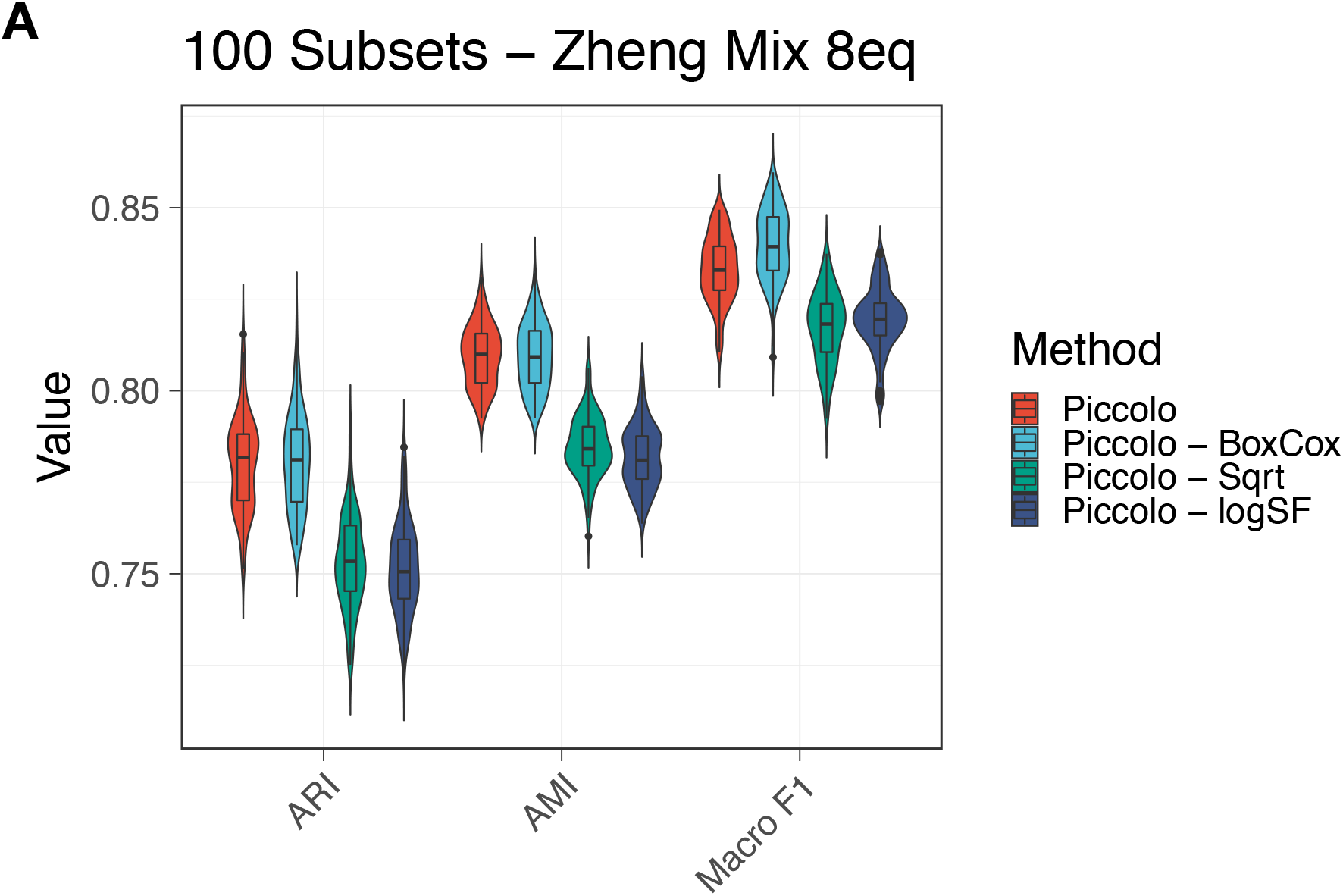
Comparison between normalizations using the variance stabilization transformations implemented in Piccolo. **A**. Violin-box plots of the clustering metrics obtained for 100 subsets of the Zheng Mix 8eq data set. The colors correspond to the respective variance stabilization transformation employed during normalization using Piccolo. The default of *log* transformation is not labeled explicitly and is simply labeled as Piccolo. For all the 100 subsets, the highest values of the metrics were observed with Piccolo (red) and Piccolo - BoxCox (blue). In fact, Piccolo - BoxCox yields higher values for the Macro F1 score compared to Piccolo (Paired Wilcoxon test (2-sided):*p* = 2.78*E* − 12).

**Fig. S17.**
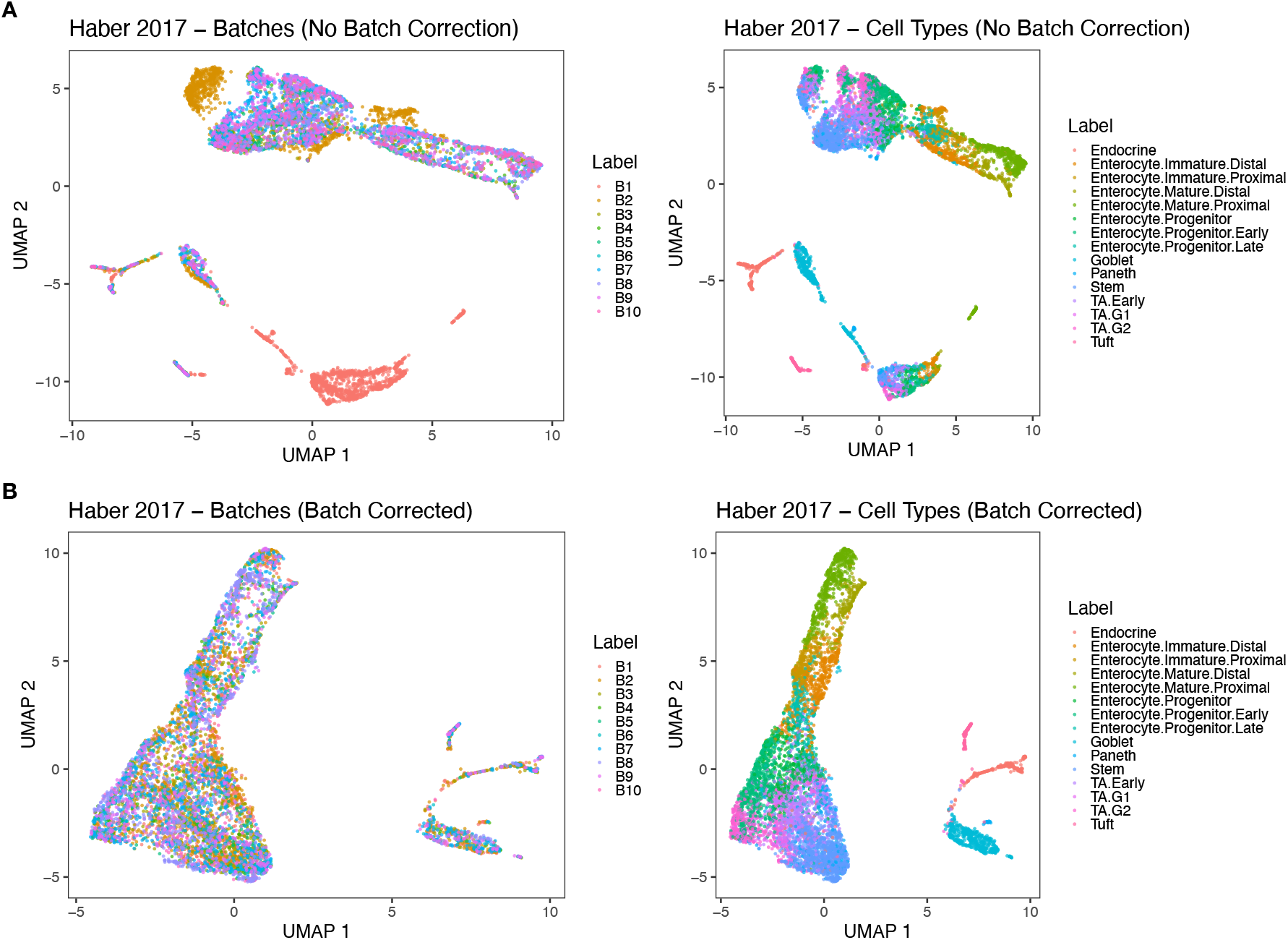
Simplistic batch effect correction enabled by Piccolo normalization works well when all or most of the biological conditions are present in all the batches. In all the plots, the dots represent cells and are colored based on the batch or cell-type labels. **A**. 2D UMAP plots after applying Piccolo without specifying batches for the Haber 2017 - Mouse Small Intestine Epithelium data set. The left panel shows cells colored by batch. We can observe clusters that clearly correspond to batches (in particular, corresponding to batches B1 (red) and B2 (golden brown)). The right panel shows the cells colored by the cell-type labels provided by the authors of the study, which reveals that within clusters belonging to different batches there are cells belonging to distinct cell-types. **B**. 2D UMAP plots after applying Piccolo wherein we specify the 10 batches. The left panel shows cells colored by batch. We now observe that cells no longer cluster based on the batches. The right panel shows cells colored by the cell-type labels provided by the authors, which clearly reveals that the cells group based on their cell-type identities after batch effect correction.

**Fig. S18.**
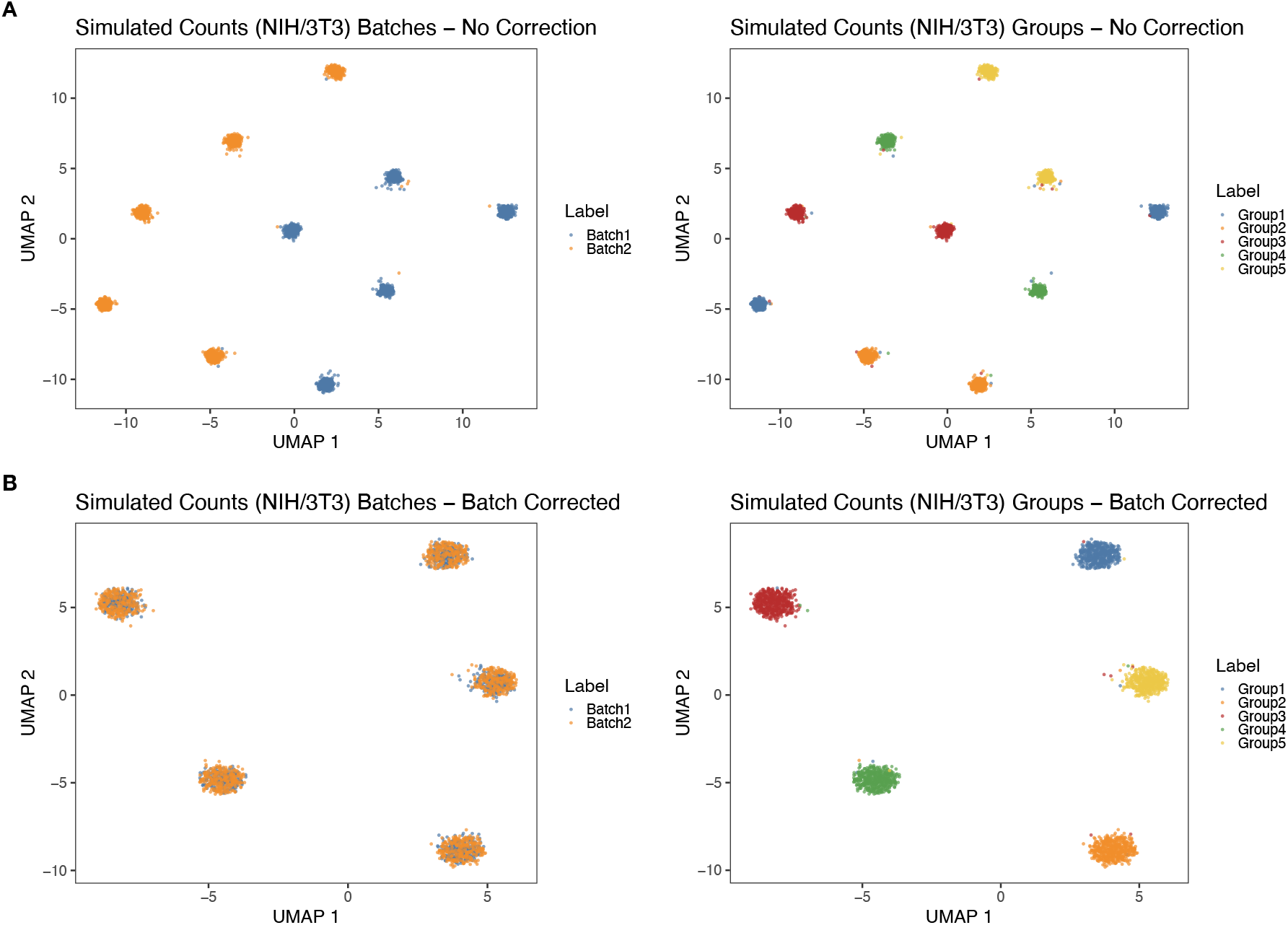
Simplistic batch effect correction enabled by Piccolo normalization works well when all the conditions (group identities) are present in all the batches. In all the plots, the dots represent cells and are colored based on the batch or group labels. **A**. 2D UMAP plots after applying Piccolo without specifying batches for a simulated counts data set in which we simulated 2 batches, with equal number of cells in each batch. The simulated counts were generated by Splat [38] using NIH/3T3 (5 groups with approximately same number of cells per group). The left panel shows cells colored by batch. We observe 10 clusters with 5 belonging to Batch1 (blue) and the other 5 belonging to Batch2 (orange). The right panel shows cells colored by the group labels, which reveals that there are 2 clusters for each group, corresponding to the 2 batches. **B**. 2D UMAP plots after applying Piccolo wherein we specify the 2 batches. The left panel shows cells colored by batch. We now observe 5 clusters, with each cluster containing cells from both batches (orange overlaid on blue). The right panel shows cells colored by the group labels, which clearly reveals that the 5 clusters correspond to the group identities of the cells.

